# Three-photon population imaging of subcortical brain regions

**DOI:** 10.1101/2025.03.21.644611

**Authors:** Hadas Frostig, Amy Monasterio, Hongjie Xia, Urvi Mishra, Baldwin Britton, John T. Giblin, Jerome Mertz, Benjamin B. Scott

**Affiliations:** Department of Psychological and Brain Sciences, Boston University, Boston, Massachusetts, USA; Graduate Program for Neuroscience, Boston University, Boston, Massachusetts, USA; Department of Biology, Boston University, Boston, Massachusetts, USA; Thorlabs Inc., Sterling, Virginia, USA; Department of Biomedical Engineering, Boston University, Boston, Massachusetts, USA

## Abstract

Recording activity from large cell populations in deep neural circuits is essential for understanding brain function. Three-photon (3P) imaging is an emerging technology that allows for imaging of structure and function in subcortical brain structures. However, increased tissue heating, as well as the low repetition rate sources inherent to 3P imaging, have limited the fields of view (FOV) to areas of ≤ 0.3 mm^***2***^. Here we present a Large Imaging Field of view Three-photon (LIFT) microscope with a FOV of ***>***3 mm^***2***^. LIFT combines high numerical aperture (NA) optimized sampling, using a custom scanning module, with deep learning-based denoising, to enable population imaging in deep brain regions. We demonstrate non-invasive calcium imaging in the mouse brain from ***>***1500 cells across CA1, the surrounding white matter, and adjacent deep layers of the cortex, and show population imaging with high signal-to-noise in the rat cortex at a depth of 1.2 mm. The LIFT microscope was built with all off-the-shelf components and allows for a flexible choice of imaging scale and rate.

## Main

Two-photon (2P) microscopy [1–4] with calcium indicators [5–7] has revolutionized in-vivo brain imaging, as it allows noninvasive recording of neural activity in scattering brain tissue. Standard 2P systems can image with subcellular resolution several hundreds of microns into the brain noninvasively, providing access to multiple layers of the mouse cortex [8, 9]. Several large field-of-view (FOV) 2P instruments have been developed and utilized to image large neuronal populations [10–16]. However scattering of the excitation light and out-of-focus signal degrade image quality in depth [3, 17, 18]. As a result, calcium imaging in the deeper layers of the cortex can be challenging with standard instruments, and subcortical structures are only accessible with invasive methods [19–21]. In larger animals, such as the rat, noninvasive 2P microscopy is typically limited to supragranular layers [22].

In recent years, three-photon (3P) microscopy has emerged as a neuroimaging technique that allows access to deeper layers of the brain [23–26], including the mouse hippocampus [27–30]. In 3P microscopy, three infrared photons are simultaneously absorbed in a molecule to elicit fluorescence at a visible wavelength. As a result, the excitation wavelength is longer than that of 2P microscopy and therefore the scattering of the excitation light is reduced [31, 32]. Furthermore, since 3P absorption is a fifth-order nonlinear effect, the signal intensity, *S*, depends on the excitation pulse energy, *E*_*p*_, as

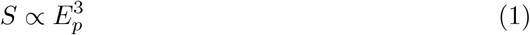

assuming a fixed pulse duration. The higher-order nonlinearity of the signal reduces out-of-focus background and improves optical sectioning compared to 2P imaging. Unfortunately, the longer excitation wavelength used in 3P also increases the absorption of the excitation light in tissue [31]. Hence 3P imaging is limited by tissue heating, which restricts the power of the excitation source.

While multiple works have demonstrated the usefulness of 3P imaging, the fields of view imaged at 1 mm depth have been limited (areas ≤ 0.3 mm^2^ [28, 29, 33]) for several reasons. First, the high pulse energies required for 3P imaging dictate low repetition rate excitation sources. This limits the rate at which data can be acquired, restricting the number of focal spots and the FOV size that can be imaged at rates compatible with calcium imaging. In principle the repetition rate could be increased, together with the excitation power, to preserve excitation pulse energies. Yet this approach results in excitation powers that are beyond the limitations generally accepted for tissue heating [32, 34].

Here we present LIFT (Large Imaging Field-of-view Three photon) microscopy, capable of recording neural activity from a ≥ 3 mm^2^ FOV at a depth of 1 mm in vivo. The LIFT microscope uses a high NA optimized sampling scheme, which we show not only enables large FOV 3P calcium imaging despite the limited data acquisition rate, but also maximizes the signal-to-noise ratio (SNR) and mitigates the challenges associated with tissue heating. We built a custom scanning module that performs optimized spatial sampling over large FOVs while allowing for maximal versatility in imaging scale and rate. Combining this optical system with deep-learning based denoising, we demonstrate a 10-fold increase in the FOV size of 3P functional imaging 1 mm deep in the mouse brain, and 15-fold increase in the number of neurons recorded from compared to previous results. We demonstrate the first large FOV 3P imaging in a rat, and image *>*900 GCaMP-labelled neurons in the rat motor cortex, at a depth of 1.2 mm, with high SNR. Our system uses all off-the-shelf components and is easily reproducible.

## Results

### Design constraints

Conventional approaches for increasing FOV, commonly used in 2P imaging and other point-scanning imaging methods such as OCT [35], are not well-suited for 3P imaging. Multiple 2P systems use larger focal spots in order to scan a larger FOV [13, 29] (Fig. 1a), since the 2P signal from a cell does not depend on the size of the focal spot, as long as the point-spread function (PSF) does not exceed the cell size [36] (black line, Fig. 1b). In 3P imaging, however, the signal depends on the focal spot diameter, *d*, as *S ∝* 1*/d*^2^, causing a rapid loss in signal with spot size increase (orange line, Fig. 1b). Another common approach for increasing the FOV is to increase the number of focal spots imaged per frame (Fig. 1c). In order to preserve the frame rate, the focal spots need to be scanned either more quickly [11, 15] or in parallel, using spatiotemporal multiplexing [29, 37–39]. While this approach results in mild signal degradation in 2P imaging (black line, Fig. 1d), it causes a rapid decay in signal in 3P imaging. In order to increase the FOV diameter *N* times in 3P imaging by scanning faster, the repetition rate must be increased by *N* ^2^, reducing the pulse energy to *E*_*p*_*/N* ^2^, and causing the signal to drop as 1*/N* ^6^ due to the cubic dependence in Eq. 1 (green line, Fig. 1d). Similarly, in order to increase the FOV diameter *N* times using spatiotemporal multiplexing, the excitation power must be split between *N* ^2^ beamlets, reducing the signal by 1*/N* ^6^. While this approach can be useful for large FOV 3P imaging in deep cortical regions [33], the large reduction in signal makes imaging subcortical regions challenging (see Supplemental Note 2).

**Fig. 1.**
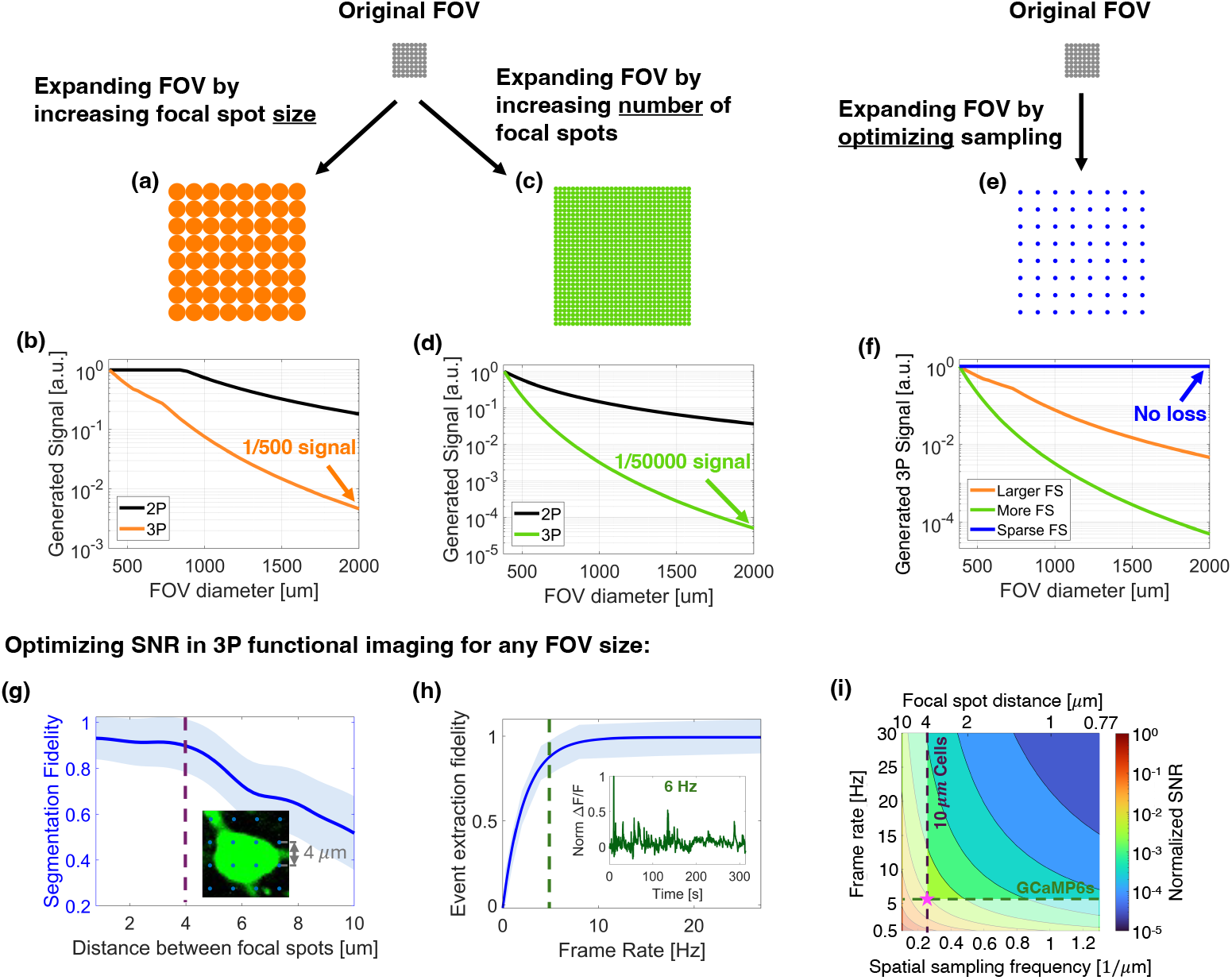
Optimized sampling provides a solution to the challenges in large FOV 3P imaging. (a)–(d) The challenges in expanding the FOV in 3P imaging. Two common methods of expanding FOV are increasing the focal spot size (panel a) and increasing the number of focal spots imaged per frame by scanning faster or by multiplexing (panel c). While these methods result in mild signal degradation in 2P imaging (black lines in b and d) and other scanning microscopy methods, they reduce the signal in 3P imaging by 2.5 (orange line in b) to 4.5 (green line in d) orders of magnitude for a 2 mm FOV diameter, respectively. (e)–(h) Expanding the FOV by increasing sparsity in space (panel e) and time preserves the amount of signal (blue line in f), the fidelity of cell segmentation (panel g) and calcium event extraction (panel h) up to 4 *µ*m focal spot spacing (inset of g) and 6 Hz frame rate (inset of h), which is the sampling scheme used in LIFT. The shaded blue region in panels g-h represents one standard deviation from the mean fidelity, see Supplemental Note 1. (i) The SNR in 3P functional imaging as a function of the frame rate and distance between focal spots, for a shotnoise limited system. Even for small FOVs, choosing the lowest frame rate compatible with indicator kinetics (GCaMP6s shown in green) and the largest focal spot spacing that allows to resolve cells (10 *µ*m cells shown in purple) will yield the optimal SNR (pink star).

### System design

The LIFT microscope was designed to accommodate the largest FOV that can be imaged with cellular resolution at imaging rates compatible with GCaMP6s kinetics, as well as to maximize the SNR of 3P functional imaging at around 1 mm imaging depth. Additionally, LIFT was designed to be a versatile system, that could perform imaging on different scales and rates, and one that could be built using only off-the-shelf components.

To simultaneously maximize both the FOV and the SNR, we designed LIFT to sample the focal plane as sparsely as possible without undersampling cells [16], and record at the minimal frame rate that allows to resolve events after denoising (Supplemental Note 1., Extended Data Fig. 1-2). Using high NA spatiotemporally optimized sampling (Fig. 1e) allows to expand the FOV while preserving the signal magnitude (Fig. 1f, blue line). Furthermore, due to the highly nonlinear nature of the 3P signal, optimizing spatial and temporal sampling maximizes the SNR of 3P calcium imaging for *any* FOV size (Fig. 1i, Supplemental Note 2). To determine the optimal focal spot spacing and frame rate, we performed measurements to quantify the the fidelity of cell segmentation for different focal spot spacings (Fig. 1g) and the fidelity of calcium event extraction for different frame rates (Fig. 1h). We chose a focal spot spacing of 4 *µ*m (purple dashed line in Fig. 1g-i, Extended Data Fig. 3) and a frame rate of 6 Hz (green dashed line in Fig. 1h-i) which allowed maximal sparsity with minor reduction in fidelity (*≥*0.9 for both cell segmentation and event extraction). In addition to maximizing SNR, distancing the focal spots reduces local tissue heating, allowing to increase the power incident on the brain surface and further increasing SNR (Supplemental Note 2, Extended Data Fig. 4).

LIFT was also designed to provide flexibility in imaging scale and rate. Versatility is required for recording activity from neural components of varying sizes (soma, axons, dendrites), accommodating indicators with different response times, as well as simply to navigate in the brain during an imaging session. All the above requirements dictate a scanner that is able to perform rapid, large-angle beam scanning with a customized scan waveform yet preserve flexibility in scan speed and FOV size. Such scanning cannot be performed with the four commonly-used scanners: galvanometer, resonant scanner, polygon scanner or microelectromechanical (MEMS) scanning mirror (see Supplemental Note 3).

To that end, we implemented a custom scanning module, as outlined in Fig. 2a. Two galvanometers are used in a relay configuration, so that their scan angles add. This design both preserves the advantages of the galvanometer, allowing to tailor the scan pattern and rate to the sample, and doubles the scan speed of a single galvanometer, producing the range of speeds necessary for large FOV 3P imaging of neural activity. While galvanomoters are typically used to image only the central portion of the scan line, we implemented imaging on turnaround (Fig. 2b, top). Imaging on the turnaround portion of the scan, defined as the portion of motion that deviates from linearity due to directional change, maximizes the FOV for a given frame rate. Fig. 2b shows a comparison of the FOV achievable with our waveform with a single galvanometer, a single galvanometer with turnaround imaging, and LIFT scope.

**Fig. 2.**
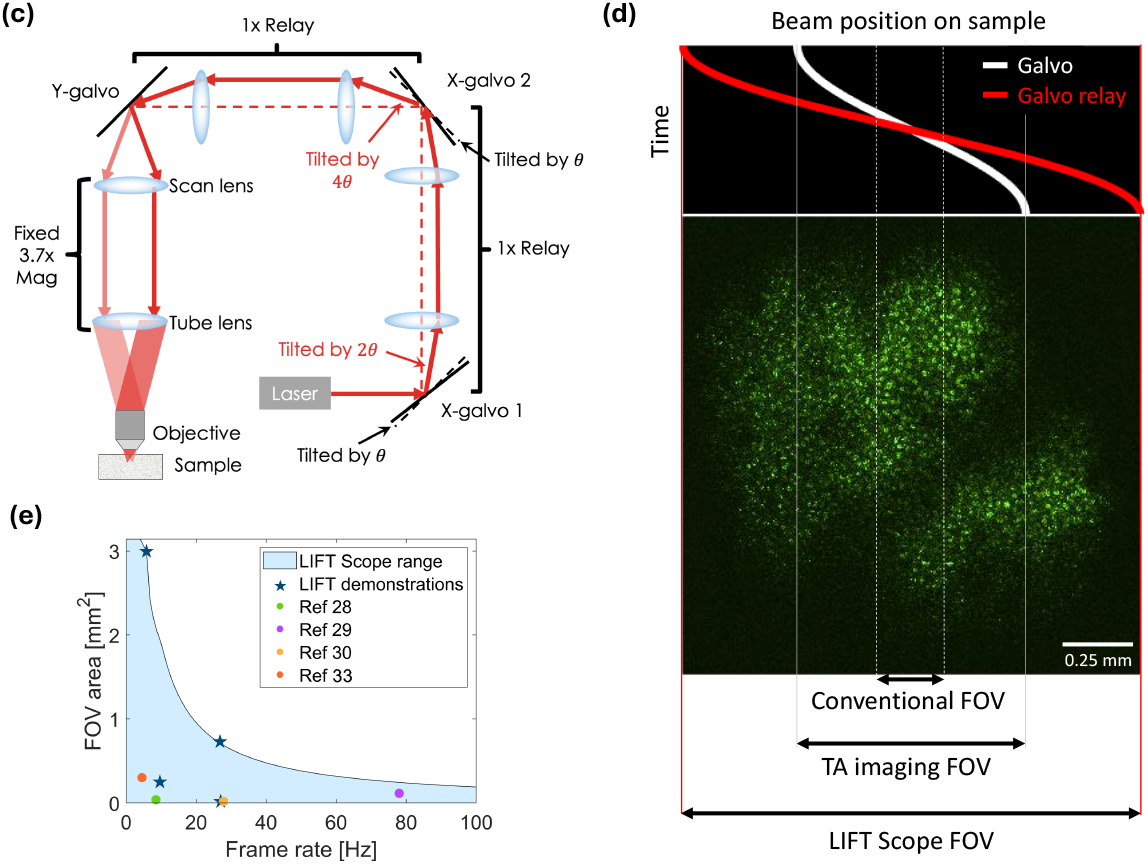
The LIFT microscope design and performance. (a) A schematic of the optical path. A 1:1 relay between X-galvanometer 1 and X-galvanometer 2 (galvo relay) is used to double the scan angle. (b) A comparison of the fields-of-view achievable at frame rates compatible with GCaMP6s kinetics with a conventional galvanometer scan (conventional FOV), a galvanometer scan with turnaround imaging (TA imaging FOV), and LIFT scope (LIFT scope FOV). The top panel shows the scanned beam position on the sample in a single line in the frame for a single galvanometer (white) and a galvanometer relay (red). The image shown is an in-vivo image of rat brain expressing Ribo-jGCaMP8m at a depth of 1.2 mm. (c) A comparison between the fields-of-view achievable at different frame rates using LIFT scope and previous 3P systems that were used to image at ∼ 1 mm depth in-vivo. The filled-in light blue area denotes the operational range of LIFT scope, and the blue stars denote LIFT imaging demonstrations shown in this work.

The performance of the LIFT microscope is depicted in Fig 2c. The area under the curve represents the operational range of LIFT scope, and combinations of FOVs and frame rates demonstrated in this work are marked with stars. Using optimized sampling, LIFT scope can image a 2 mm diameter FOV at a frame rate of 4.7 Hz, and a 1.65 mm x 0.45 mm FOV at a rate of 27 Hz. On the other hand, in applications that call for finer spatial sampling, LIFT scope can provide <0.5 *µ*m lateral sampling with FOVs that are 2 - 5 times larger than conventional 3P systems [28].

In order to resolve activity from large populations within the FOV, we denoised the functional imaging data with a self-supervised denoising algorithm, DeepCAD-rt [40, 41]. We studied the performance of DeepCAD-rt in comparison with high-performance filtering techniques, and showed that DeepCAD-rt significantly enhances SNR and hence the number of cells with resolvable activity, while preserving fluorescence traces (Extended Data Fig. 5 and 6).

While many large FOV microscopes employ large, custom-designed optics, LIFT scope was built entirely from off-the-shelf, readily-available components (see Methods). Since the repetition rate of the excitation light (1 MHz has been shown to be optimal for imaging at 1 mm depth [42]) constrains the number of focal spots that can be imaged at frame rates suitable for calcium imaging, the scanned FOV is limited to 2 mm x 2 mm for 4 *µ*m sampling at approximately 5 Hz frame rate regardless of microscope design. For this FOV, a commercial objective exists that also offers the high NA required for 3P imaging (Nikon 16x, 0.8 NA). By carefully designing our system to match this objective and our scanner to match off-the-shelf scan lenses, we were able to avoid bulky, high-cost custom optical components.

### Multiscale imaging of structure and function in mice

To validate the high resolution structural imaging capabilities of LIFT scope in vivo, we performed large FOV volumetric (2 x 2 x 1.23 mm) imaging of GCaMP6s-labeled cells in mice with 1 *µ*m lateral sampling (Fig. 3a and Supplementary video 1). Processes and neurons were visible in the cortex (Fig. 3a and 3b) as well as in the dorsal region of CA1 (Fig. 3a and 3c), and the intermediate white matter appeared as a lower cell density region (which appears dark in Fig. 3a, curving down to the left). Since our FOV is larger than the horizontal cross-section of the hippocampus, regions of the white matter surrounding it were visible, as well as adjacent regions of deep cortex (Supplementary Video 1). At depths *>* 1.13 mm, we observed a change in the structure of the cell layer, which may indicate a transition into CA2.

**Fig. 3.**
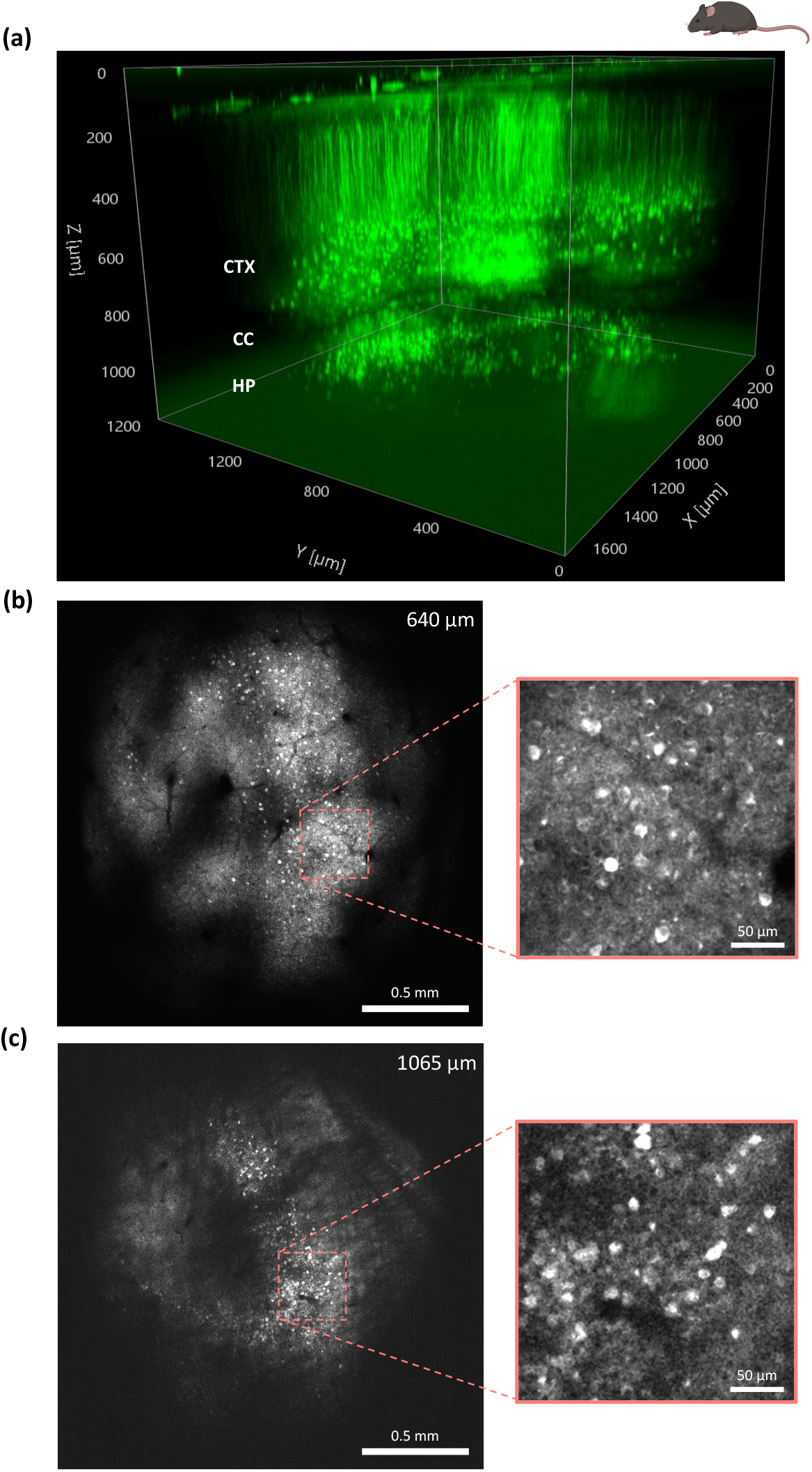
Large FOV high resolution structural 3P imaging in a mouse expressing GCaMP6s. (a) 3D rendering of a structural stack recorded with 1 *µ*m lateral sampling, at a depth of 0 - 1.2 mm below dura. The cortex (CTX), corpus collasum (CC), and hippocampus (HP) are all visible. Representative images from the stack, recorded at (b) 640 *µ*m depth, corresponding to layer 5/6 in the cortex, and (c) 1065 *µ*m depth, corresponding to hippocampus and surrounding regions. Insets: magnified views.

We then used LIFT to record calcium activity from a large FOV in CA1 and adjacent regions using optimized spatiotemporal sampling. We performed calcium imaging in head-fixed mice running on a horizontal disk treadmill (Fig. 4a). The recorded datasets were denoised with deepCAD-rt, which nearly tripled the number of cells detected, improved cell segmentation fidelity, and improved calcium event inference across the field of view (Extended Data Fig. 5 and 6), compared to conventional filters (Kalman and running average [33]). Automated region-of-interest (ROI) extraction from the denoised data recorded at a depth of 981 *µ*m revealed *>*1500 active cells in this FOV (Fig. 4b-c, Supplementary video 2). A high density of active cells was seen in CA1 (labeled HP), as well as in the adjacent somatosensory cortex (labeled CTX), whereas a lower cell density was apparent in the intermediate white matter in the corpus callosum (labeled CC), in accordance with literature [28, 43] (Fig. 4b; see Fig. 5a for structure). The majority of extracted activity traces had 3 < SNR < 8 (Fig. 4d; see methods for SNR definition). Representative traces from various SNR levels showed clear fluorescence transients even for the lower SNR groups (Fig. 4e). The cell segmentation procedure showed high fidelity, with an F-score of 0.9 (see methods and Supplementary Note 1). Next, we used LIFT to record activity throughout the volume of CA1, at depths ranging from 930 *µ*m to 1081 *µ*m, with an FOV of 3 mm^2^ (Fig. 5). LIFT’s large FOV enabled recording activity from multiple layers in CA1 (stratum oriens, stratum pyramidale and stratum radiatum) (Fig. 5i-n), as well as from the adjacent corpus callosum and somatosensory cortex (Fig. 5c-f), simultaneously. Due to the size of the FOV, the ring-shaped cross-section of SP and SO is visible in the data (Fig. 5i and 5l). Finally, we validated LIFT’s ability to correlate neural activity to behavior, by examining calcium activity at 1 mm depth during locomotion. The data revealed a distinct cell population in deep motor cortex that showed elevated activity during locomotion bouts (Fig. 4f).

**Fig. 4.**
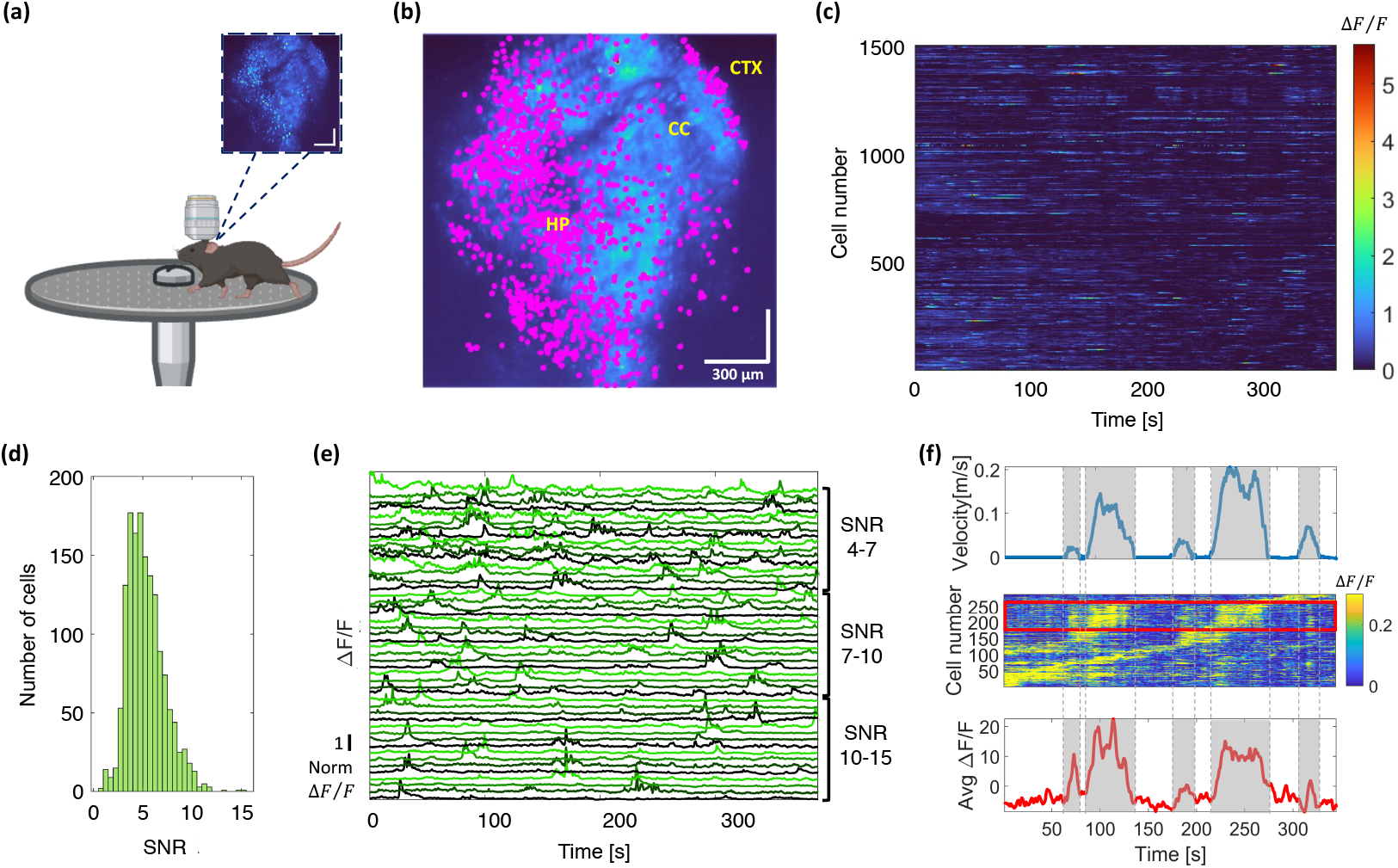
3P subcortical population imaging of neural activity in awake, behaving mice. (a) Schematic of approach. 3P calcium imaging was performed in head-fixed mice running on a horizontal disk treadmill. Inset: the MIP of a large FOV time series recorded at a depth of 981 *µ*m in the mouse brain. (b) Contours of 1540 active cells, plotted in Magenta, on top of the MIP. Active cells from the CA1 subfield of the hippocampus (HP), adjacent corpus collasum (CC), and cortex (CTX) are visible in the data (see Fig. 5a). (c) Calcium dynamics from all 1540 active cells. (d) A histogram of the SNRs of the traces shows that the majority have 3 < SNR < 8. (e) Examples of traces from various SNR ranges. Clear transients are visible even for the lowest SNR group. (f) The correlation between locomotion and calcium activity recorded at a depth of 1 mm in mouse motor cortex. Top: The recorded treadmill velocity. Middle: The extracted traces sorted by the time of maximum correlation with treadmill velocity. The red box indicated traces of cells whose activity is correlated with locomotion. Bottom: The averaged 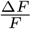 of the traces in the red box. Increased activity is observed during locomotion bouts (gray background). Imaging was performed with 4 *µ*m lateral sampling and 6 Hz frame rate.

**Fig. 5.**
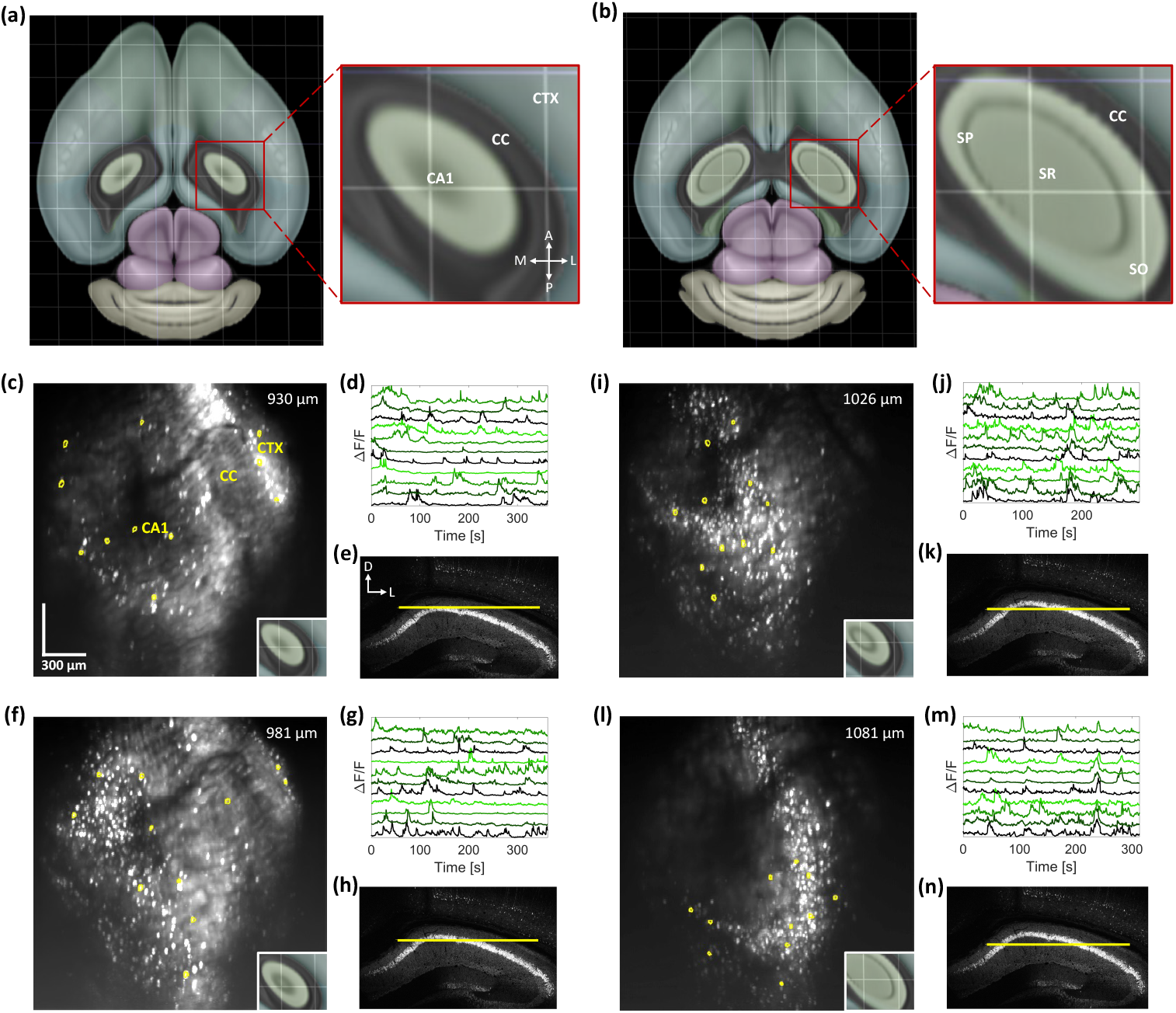
Neuronal activity recordings in awake, behaving mice from large cross sections through multiple layers of CA1 and adjacent brain regions. (a) A horizontal section through the mouse brain at the top of CA1, showing the geometry of CA1 and the surrounding corpus callosum (CC) and cortex (CTX). Grid size, 1 mm. White arrows indicate anterior (A), posterior (P), medial (M), and lateral (L) directions. (b) A horizontal section deeper into CA1, showing the geometry of the layers in CA1: stratum oriens (SR), stratum pyramidale (SP) and stratum radiatum (SR) (a and b reproduced from the Allen brain explorer 2 [44]). (c) The maximum intensity projection (MIP) of the data recorded at z = 930 *µ*m with examples of ROIs plotted in yellow on top. Inset: a zoom in on a horizontal section from roughly the same depth, showing the approximate geometry of the imaged FOV. (d) the corresponding activity traces from those ROIs. (e) A histology slice from the same brain, with the approximate imaging area marked (yellow line). White arrows indicate dorsal (D) and lateral (L) directions. (f)-(h) Same as c-e but for z = 981 *µ*m. (i)-(k) Same as c-e but for z = 1026 *µ*m. (l)-(n) Same as c-e but for z = 1081 *µ*m. The ring-shaped cross-section of SP and SO is visible in panels i and l. Imaging was performed with 4 *µ*m lateral sampling and a 6 Hz frame rate.

To evaluate the versatility of the LIFT scope, we tested its capability to record neuronal activity at faster frame rates and higher spatial resolutions. A dataset acquired at a 27 Hz frame rate, covering a 0.73 mm^2^ FOV in the CA1 region of the mouse hippocampus, yielded high SNR traces from 400 active cells (Extended Data Fig. 7). Similarly, datasets acquired with lateral sampling between 0.8 and 2 µm in the mouse hippocampus demonstrated consistently high SNR. An example data set, acquired with 2 *µ*m lateral sampling and an 11 Hz frame rate, which was not denoised or filtered, is shown in Extended Data Fig. 8 (for 0.8 *µ*m lateral sampling see Extended Data Fig. 1). Together, these datasets indicate the flexibility of LIFT and its ability to perform high SNR 3P imaging at multiples scales and rates.

### High SNR large FOV neuronal imaging in deep rat brain

Larger brained animals, such as rats and non-human primates, are of interest in neuroscience research due to their complex behavioral repertoire and physiological similarity to humans. However, the brains of these species contain thicker cortical layers and lower neuronal density compared to mice. [22, 45, 46]. Thus recording activity from large populations of neurons requires large FOV imaging with considerable depth access.

To evaluate the performance of LIFT in a larger brained model, we performed large FOV high resolution structural imaging of Ribo-jGCaMP8m-labeled cells [47, 48] in the secondary motor cortex (M2) of rats (Fig. 6). We used a soma-targeted indicator to mitigate the effects of the higher neuropil-to-soma ratio in the rat brain [22]. The results revealed large cell populations (n = 700 - 1400, depending on the depth of imaging relative to the location of the cell layer) throughout the cortical column (Fig. 6g). At the largest depth, 1200 *µ*m, over 900 cells were visible, and the majority of SNRs were between 4 and 14 (Fig. 6h, see methods for SNR definition). Histological images confirmed labeling through the entire depth of M2 (Fig. 6i). Our imaging depth was not limited by SNR but by the geometry of our cranial window (see methods).

**Fig. 6.**
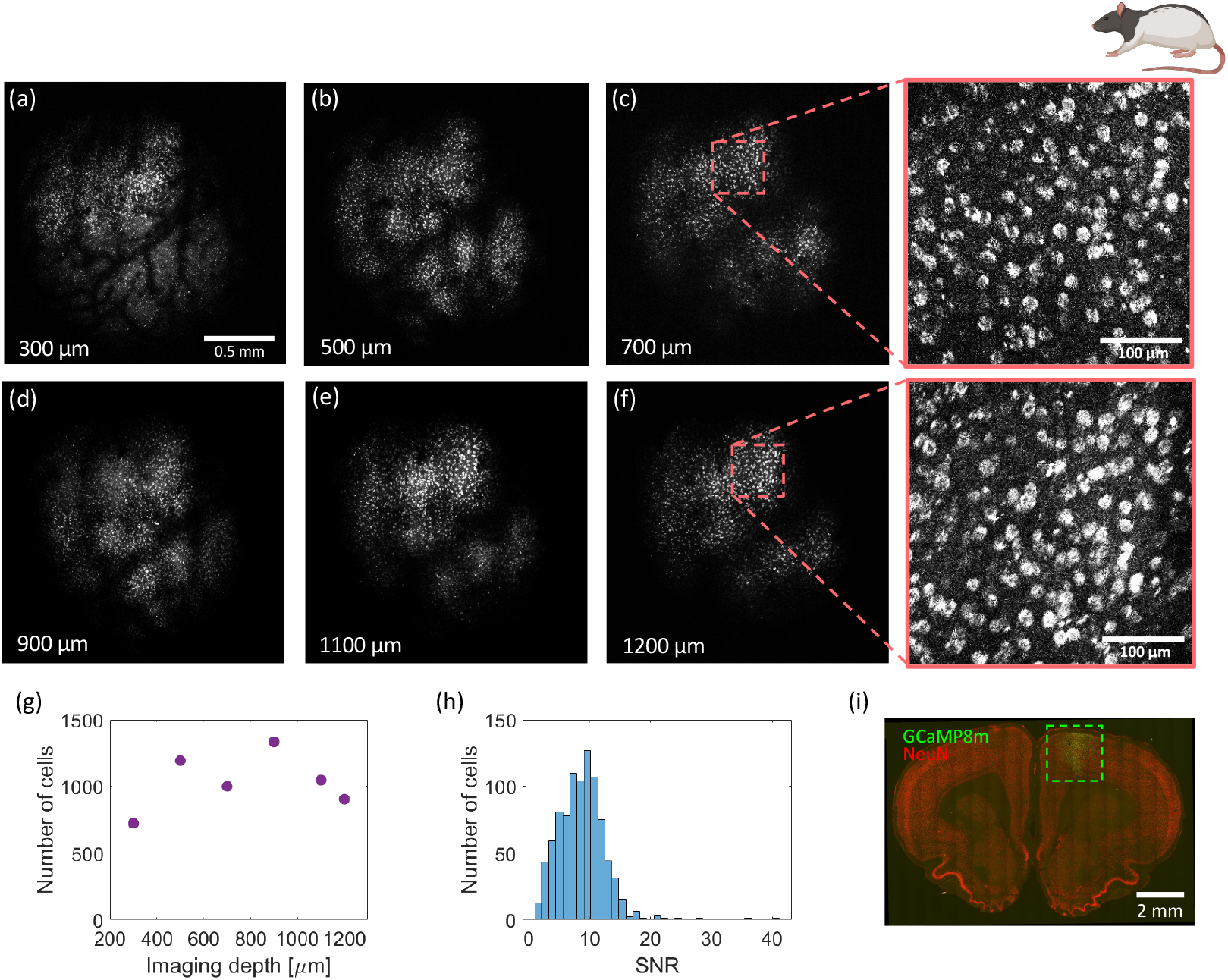
Large FOV 3P population imaging in a rat expressing Ribo-jGCaMP8m. Images taken at depths of (a) 300 *µ*m (b) 500 *µ*m (c) 700 *µ*m (d) 900 *µ*m (e) 1100 *µ*m and (f) 1200 *µ*m below dura, with 1 *µ*m lateral sampling. No post processing was performed on the images. Insets: magnified views of c and f. (g) The number of cells segmented in each image. (h) The SNR histogram of the cells segmented in the 1200 *µ*m depth image. Imaging was performed in an anesthetized rat. (i) Confocal images of a histological slice from a brain of an example rat. Green channel: Ribo-jGCaMP8m, Red Channel: NeuN. The green box indicates the area of GCaMP expression.

## Discussion

In order for 3P imaging to mature into a technology that allows researchers to study questions about brain function and dynamics, systems capable of recording activity from large cell populations and large areas are necessary [49, 50]. Many previous findings have shown long correlation distances of activity patterns even for simple sensory inputs [51–53], showing that probing small, local neuronal populations may not provide the full picture necessary for neuroscience research.

Using LIFT scope, we demonstrated large FOV 3P imaging of structure and function, as deep as 1 - 1.2 mm below cortical surface, in two model rodent species. Additionally, LIFT scope enabled functional imaging from a large population of neurons (*>*1500), and several adjacent brain regions simultaneously. The ability to record activity from large fields of view in deep brain opens the door to the study of large-scale neuronal dynamics in brain regions that were inaccessible previously with noninvasive methods. In smaller animals, the large FOV can serve to record from adjacent functionally-related deep brain regions, such as the different layers of CA1, while in larger animals with lower neuronal densities, the FOV size can be exploited to record activity from large cell populations in a single deep region, such as layer 5/6 in the rat. In both of these applications, invasive methods would cause damage to the brain area studied and could severely compromise the results of the study.

In the present work, the imaging depth and/or SNR were limited by the available post-objective power. This is due to the low throughput of our optical path from the source to the sample, with 56% of the power lost on the objective itself, since it is designed for wavelengths ≤ 1100 nm. As more off-the-shelf objectives designed for 1300 nm light become commercially available, this limitation should be eliminated in the near future. Since optimized sampling alleviates the restrictions imposed by tissue heating (Supplemental Note 2, Extended Data Fig. 4), higher post-objective powers could potentially allow to increase the repetition rate while preserving excitation pulse energy. Thus more data points could be acquired at the same frame rate, enabling the expansion of the FOV beyond that demonstrated in this work. Furthermore, since our scanner is based on galvanometers, the ability to use scan patterns other than raster scanning is preserved, which can be exploited to increase the frame rate or FOV as well [54], or to perform targeted-illumination microscopy [55].

## Methods

### Excitation Source

The excitation source was a wavelength-tunable optical parametric amplifier (Opera- F, Coherent), pumped by a femtosecond amplifier (Monaco, Coherent). The Opera-F was operated at a 1300 nm wavelength and a 1 MHz repetition rate, and produced an output power of up to 1.8 W with ∼ 50 fsec pulses. Pulse width was measured with an autocorrelator (FSAC, Thorlabs) which measured predominantly third-harmonic signal and therefore the pulse width was calculated using the formula for a thirdorder autocorrelation trace. Pulse dispersion from the optics within the laser, as well as from the optics of the microscope, was compensated using a home-built singleprism compressor [56]. The excitation power was controlled using a half-waveplate on a motorized rotation stage and a polarizing beamsplitter cube.

### LIFT microscope setup

The excitation light was point-scanned over the sample using our custom-built scanner (Fig. 2a). The scanner is composed of three high-performance galvanometers (Saturn 5, ScannerMax) conjugated to one another in a ‘4f’ configuration using two pairs of identical wide FOV scan lenses (LSM54-1310, Thorlabs). The first two galvanometers scan the beam in the same direction (the fast axis) so that their scan angle adds up, and the third galvo scans in the perpendicular direction (the slow axis). The beam was expanded to fill the back aperture of the objective using another scan lens (LSM54- 1310, Thorlabs) and a tube lens (TL200-3P, Thorlabs). We used a water-dipping objective with 0.8 NA and a 3 mm working distance (CFI75 LWD 16X W, Nikon). The imaging subject’s transverse position was varied with a motorized stage (PLS-XY, Thorlabs) and its axial position was varied by scanning the position of the objective using another motorized stage. The signal was epicollected through the objective and then separated from the excitation light by a dichroic beam splitter (FF665-Di02- 25×36, Semrock). To block the third-harmonic and other background signals from the sample, we used a band pass filter (520/70BP, 87-749, Edmund) before detection. The 3P signal was detected with a hybrid photo detector (R11322U-40, Hamamatsu), and the current was converted to voltage and amplified using a transimpedance amplifier (HCA-400M-5K-C, Femto). Analog-to-digital conversion was performed by a data acquisition card (ThorDAQ, Thorlabs) and the sampling was gated (∼ 1 ns gate width) and triggered by a reference signal from the laser. ThorImageLS 4.3 (Thorlabs) was used to control image acquisition and was custom modified to enable imaging on the turnaround portion of the galvanometer motion.

### Data acquisition and analysis

The data acquisition parameters for all figures are listed in Extended Data Table 1.

### Mouse

Mouse structural images (Fig. 3) were acquired with 1 *µ*m lateral sampling, a 4 *µ*s dwell time, and a 155 *µ*s turnaround time. Images were averaged over 5 frames between 700 - 1230 *µ*m depth, and were not averaged between 0 - 700 *µ*m depth. To generate Fig. 3a, the images in the volume stack were registered using the StackReg plugin in Fiji, and 3D rendering was performed with Imaris. No postprocessing was used for Fig. 3b and c. Mouse neuroactivity (Fig. 4 and 5) was recorded with a 1 *µ*s dwell time, an average focal spot spacing of 4 *µ*m, and a turnaround time of 300 *µ*s (X-galvo waveform shown in Extended Data Fig. 3). The excitation power on the brain surface was 10 - 160 mW. The raw time series was denoised with DeepCAD-RT by first training a model on the dataset and then denoising the dataset with that model. Both training and denoising were done with 10 epochs, an overlap factor of 0.25, an xy patch of 150 (for the 3 mm^2^ FOV data) or 50 (for smaller FOV data), and a t patch of 150 (for longer movies) or 50 for (shorter movies). The denoised time series was unwarpped to correct for the nonlinearity of the galvanometer motion. The resulting time series was then Gaussian filtered to remove other galvanometer-related artifacts, and motion corrected with the rigid version of NoRMCorre [57]. Initial cell segmentation was performed with CNMF using the elliptical search method [16] and a minimum signal-to-noise parameter of 3. The ROIs were then filtered to remove highly-correlated components (correlation coefficient *>* 0.85), eliminating over-segmented cells. ROIs outside of the optical FOV were eliminated as well. Traces presented in Fig. 4 and 5 were normalized to their maximum value. SNRs were calculated by subtracting the mean value of the trace from the maximum value of the trace, and dividing by the standard deviation of the trace values. The fidelity of cell segmentation was assessed in two ways. First, by comparing the extracted ROI set from 4 *µ*m sampling data to the hand-annotated ground truth. Second, by recording high resolution data and downsampling it to yield 4 *µ*m sampling as detailed in Supplemental note 1. The F-score was computed from 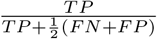 where TP, FN and FP are the true positive rate, false negative rate and false positive rate, accordingly. The F-score calculated using both methods was 0.9.

### Rat

Rat structural images (Fig. 6) were acquired with 1 *µ*m lateral sampling and a 4 *µ*s dwell time. The excitation power on the brain surface was 50 - 120 mW. The images presented in Fig. 6 were averaged over 5 frames between 900 - 1100 *µ*m depth, over 3 for 300 *µ*m depth and otherwise over 10. No post processing was performed. Cell segmentation was performed using Cellpose 2.0 [59] with manual correction. To estimate the SNR of cell imaging, the noise was taken to be the standard deviation of the pixel values in areas that do not contain visible cells. To accommodate for the ring shape created by nuclear exclusion, the signal was taken to be the mean value of the top 30% of pixels in a segmented cell body.

### Animal imaging protocols and surgical procedures

All animal use procedures were approved by the Boston University Institutional Animal Care and Use Committee (protocol numbers 201900084, 201800577).

### Mouse

C56BL6/J mice (Charles River #027) were group housed with littermates prior to experiment start and given food and water *ad libitum*. Prior to imaging experiments, mice were habituated for 5 days to head-fixation. For all surgeries, mice were initially anesthetized with 3.0% isoflurane inhalation during induction and maintained at 1.5%–2% isoflurane inhalation through stereotaxic nose-cone delivery (oxygen 1L/min). Ophthalmic ointment (Systane) was applied to the eyes to provide adequate lubrication and prevent corneal desiccation. 2.0% lidocaine hydrochloride was topically applied as local analgesia prior to midsagittal incision, using an 11 blade scalpel, of the scalp skin to expose the skull. A 0.1 mg/kg intraperitoneal (IP) dose of buprenorphine was administered at the beginning of surgery. Neurons were labeled using AAV encoding GCaMP6s in wild-type mice. 100 - 200 nL of GCaMP6s were injected at 4 sites (1.7 ML, -2.0 AP, -1.6 DV, -1.2 DV, -0.8 DV and -0.4 DV for CA1 imaging, and 1.2 ML, -1.08 AP, -1.6 DV, -1.2 DV, -0.8 DV and -0.4 DV for deep motor cortex imaging). Injections were delivered with either a 10 *µ*L Hamilton syringe with an attached 33-gauge beveled needle, or a pulled borosilicate capillary tube, at a rate of 100 nL/min (Nanoject, Drummond Scientific). For the window implant, a 3 mm diameter, 0.5 mm deep stainless steel circular cannula (Ziggy’s Tubes & Wires) was affixed to a 3 mm glass coverslip (#1 thickness, Thomas Scientific 1217N66) using UV curable optical adhesive (Norland). The cannula window was lowered carefully into the craniotomy site and Vetbond was used to adhere the edges of the cannula to the skull. The scalpel and drillbit were used to hatch the skull surface for best adhesion to the headplate. A custom stainless steel kinematic headplate [60] was affixed to the skull with the cannula centered using Metabond powder, liquid adhesive and catalyst (Parkell). Following surgery, mice were injected with a 5 mg/kg intraperitoneal (IP) dose of ketoprofen. Histological assessment verified the spread of dorsal CA1 viral targeting. Mice were administered post-operative doses of buprenorphine and ketofen for 48 hours post surgery. Mice were allowed to recover at least 1 week prior to handling/head-fixation. Imaging took place 2-4 weeks after surgery. During activity recording, mice were awake and allowed to run on a custom horizontal disk. The rotating acrylic disk was 7 inch in diameter and a soft rubber adhesive was used to provide the mice traction during running. The center of the disk was screwed into an adjustable collar (McMaster Carr 9410T1) and mounted on a rotary encoder (US Digital H1-5000-IE-D) that enabled disk position readout. An Arduino (Arduino Uno Rev 3) was used to digitize the position output and align it with image acquisition. During structural imaging, mice were anesthetized using 1.5% isoflurane and body heat was maintained using heated pads.

### Rat

Rats were anesthetized with isoflurane (5% induction, 1-2% maintenance) during the stereotaxic surgery. A 3.5 mm diameter cranial hole was drilled over the secondary motor cortex, centered at AP+ 2.25, ML +1.75. The dura was then carefully removed using 30 gauge needle tips without damaging any blood vessels. Neurons were labeled using AAV encoding Ribo-jGCaMP8m [48] which provided somatic localization of the calcium indicator. AAV9-RiboL1-jGCaMP8m (4.03x10^14^ GC/ml; viruses were made in Boston’s children hospital viral core, service request #VC-HX-4350; Addgene, plasmid #167574) was diluted 1:5 using sterile PBS. A total of 1.3 *µ*l diluted virus was delivered every 100 *µ*m between depth 1300 *µ*m and 0.1 *µ*m using a nanoject, at a rate of 100 nl/min. After virus injection, an optical implant of thickness 1.5 mm and a diameter of 3.5 mm, was placed on the surface of the cortex. The window was then sealed to the skull using vetabond and metabond. Post-operative drugs, 5 mg/kg ketoprofen and 0.02 mg/kg buprenorphine, were given to rats for two consecutive days. Rats were imaged during the fifth week post-injection, in a stereotaxic frame mounted on the microscope stage. On the day of imaging, rats were anesthetized with 80 mg/kg ketamine and 8 mg/kg xylazine. To extend the anesthesia during imaging, an additional 40 mg/kg of ketamine was used. Imaging depth was limited by the thickness of the metabond layer, which extended above the optical implant and prohibited objective-brain distances smaller than 1.25 mm.

## Acknowledgements

The authors would like to thank Dr. Anderson Chen (Neurophotonics Center, Boston University) for help with the mechanical design of LIFT scope, and Prof. David Boas (Boston University) for his advice and support. The authors also thank Eric Lieser, Hongzhou Ma, and Jeff Brooker (Thorlabs Inc.) for helpful discussions, technical assistance, and providing critical parts of the microscope. Research reported in this publication was supported by awards R01 NS116139 and R56 MH132732 from the National Institutes of Health (NIH), by the Neurophotonics center in Boston University, and by the Research Corporation for Science Advancement. A.M. is supported by the NIH DSPAN award, and H.F. is supported by the Israeli National Postdoctoral Award and by the American Committee for Weizmann Institute of Science.

## Author Contributions

J.M. and B.B.S. jointly supervised this work. H.F., J.M. and B.B.S. conceived the study. H.F. and J.M. designed the instrument with input from B.B.S.. H.F. built the instrument. A.M., H.X. and U.M. prepared the animals. H.F., A.M., H.X., U.M. and J.T.G. performed the experiments. B.B. implemented software features for the instrument. H.F., A.M., H.X. and J.T.G. analyzed the data. All authors contributed to manuscript preparation.

## Competing interests

The authors declare no competing financial interests.

## Notes

### Competing Interest Statement

The authors have declared no competing interest.

## References

[1] Denk, W., Strickler, J.H., Webb, W.W.: Two-photon laser scanning fluorescence microscopy. Science 248(4951), 73–76 (1990)

[2] So, P.T., Dong, C.Y., Masters, B.R., Berland, K.M.: Two-photon excitation fluorescence microscopy. Annual review of biomedical engineering 2(1), 399–429 (2000)

[3] Helmchen, F., Denk, W.: Deep tissue two-photon microscopy. Nature methods 2(12), 932–940 (2005)

[4] Svoboda, K., Yasuda, R.: Principles of two-photon excitation microscopy and its applications to neuroscience. Neuron 50(6), 823–839 (2006)

[5] Miyawaki, A., Llopis, J., Heim, R., McCaffery, J.M., Adams, J.A., Ikura, M., Tsien, R.Y.: Fluorescent indicators for ca2+ based on green fluorescent proteins and calmodulin. Nature 388(6645), 882–887 (1997)

[6] Nakai, J., Ohkura, M., Imoto, K.: A high signal-to-noise ca2+ probe composed of a single green fluorescent protein. Nature biotechnology 19(2), 137–141 (2001)

[7] Chen, T.-W., Wardill, T.J., Sun, Y., Pulver, S.R., Renninger, S.L., Baohan, A., Schreiter, E.R., Kerr, R.A., Orger, M.B., Jayaraman, V., et al.: Ultrasensitive fluorescent proteins for imaging neuronal activity. Nature 499(7458), 295–300 15 (2013)

[8] Helmchen, F., Svoboda, K., Denk, W., Tank, D.W.: In vivo dendritic calcium dynamics in deep-layer cortical pyramidal neurons. Nature neuroscience 2(11), 989–996 (1999)

[9] Mittmann, W., Wallace, D.J., Czubayko, U., Herb, J.T., Schaefer, A.T., Looger, L.L., Denk, W., Kerr, J.N.: Two-photon calcium imaging of evoked activity from l5 somatosensory neurons in vivo. Nature neuroscience 14(8), 1089–1093 (2011)

[10] Tsai, P.S., Mateo, C., Field, J.J., Schaffer, C.B., Anderson, M.E., Kleinfeld, D.: Ultra-large field-of-view two-photon microscopy. Optics express 23(11), 13833–13847 (2015)

[11] Sofroniew, N.J., Flickinger, D., King, J., Svoboda, K.: A large field of view two-photon mesoscope with subcellular resolution for in vivo imaging. elife 5, 14472 (2016)

[12] Chen, J.L., Voigt, F.F., Javadzadeh, M., Krueppel, R., Helmchen, F.: Long-range population dynamics of anatomically defined neocortical networks. Elife 5, 14679 (2016)

[13] Bumstead, J.R., Park, J.J., Rosen, I.A., Kraft, A.W., Wright, P.W., Reisman, M.D., Côté, D.C., Culver, J.P.: Designing a large field-of-view two-photon microscope using optical invariant analysis. Neurophotonics 5(2), 025001–025001 (2018)

[14] Lu, R., Liang, Y., Meng, G., Zhou, P., Svoboda, K., Paninski, L., Ji, N.: Rapid mesoscale volumetric imaging of neural activity with synaptic resolution. Nature methods 17(3), 291–294 (2020)

[15] Yu, C.-H., Stirman, J.N., Yu, Y., Hira, R., Smith, S.L.: Diesel2p mesoscope with dual independent scan engines for flexible capture of dynamics in distributed neural circuitry. Nature communications 12(1), 6639 (2021)

[16] Demas, J., Manley, J., Tejera, F., Barber, K., Kim, H., Traub, F.M., Chen, B., Vaziri, A.: High-speed, cortex-wide volumetric recording of neuroactivity at cellular resolution using light beads microscopy. Nature Methods 18(9), 1103–1111 (2021)

[17] Oheim, M., Beaurepaire, E., Chaigneau, E., Mertz, J., Charpak, S.: Two-photon microscopy in brain tissue: parameters influencing the imaging depth. Journal of neuroscience methods 111(1), 29–37 (2001)

[18] Theer, P., Denk, W.: On the fundamental imaging-depth limit in two-photon microscopy. JOSA A 23(12), 3139–3149 (2006)

[19] Levene, M.J., Dombeck, D.A., Kasischke, K.A., Molloy, R.P., Webb, W.W.: In vivo multiphoton microscopy of deep brain tissue. Journal of neurophysiology 91(4), 1908–1912 (2004)

[20] Dombeck, D.A., Harvey, C.D., Tian, L., Looger, L.L., Tank, D.W.: Functional imaging of hippocampal place cells at cellular resolution during virtual navigation. Nature neuroscience 13(11), 1433–1440 (2010)

[21] Ziv, Y., Burns, L.D., Cocker, E.D., Hamel, E.O., Ghosh, K.K., Kitch, L.J., Gamal, A.E., Schnitzer, M.J.: Long-term dynamics of ca1 hippocampal place codes. Nature neuroscience 16(3), 264–266 (2013)

[22] Kim, S.J., Affan, R.O., Frostig, H., Scott, B.B., Alexander, A.S.: Advances in cellular resolution microscopy for brain imaging in rats. Neurophotonics 10(4), 044304–044304 (2023)

[23] Rodríguez, C., Liang, Y., Lu, R., Ji, N.: Three-photon fluorescence microscopy with an axially elongated bessel focus. Optics letters 43(8), 1914–1917 (2018)

[24] Yildirim, M., Sugihara, H., So, P.T., Sur, M.: Functional imaging of visual cortical layers and subplate in awake mice with optimized three-photon microscopy. Nature communications 10(1), 177 (2019)

[25] Klioutchnikov, A., Wallace, D.J., Sawinski, J., Voit, K.-M., Groemping, Y., Kerr, J.N.: A three-photon head-mounted microscope for imaging all layers of visual cortex in freely moving mice. Nature methods 20(4), 610–616 (2023)

[26] Yu, C.-H., Yu, Y., Adsit, L.M., Chang, J.T., Barchini, J., Moberly, A.H., Benisty, H., Kim, J., Young, B.K., Heng, K., et al.: The cousa objective: a long-working distance air objective for multiphoton imaging in vivo. Nature Methods 21(1), 132–141 (2024)

[27] Horton, N.G., Wang, K., Kobat, D., Clark, C.G., Wise, F.W., Schaffer, C.B., Xu, C.: In vivo three-photon microscopy of subcortical structures within an intact mouse brain. Nature photonics 7(3), 205–209 (2013)

[28] Ouzounov, D.G., Wang, T., Wang, M., Feng, D.D., Horton, N.G., Cruz-Hernández, J.C., Cheng, Y.-T., Reimer, J., Tolias, A.S., Nishimura, N., et al.: In vivo three-photon imaging of activity of gcamp6-labeled neurons deep in intact mouse brain. Nature methods 14(4), 388–390 (2017)

[29] Weisenburger, S., Tejera, F., Demas, J., Chen, B., Manley, J., Sparks, F.T., Traub, F.M., Daigle, T., Zeng, H., Losonczy, A., et al.: Volumetric ca2+ imaging in the mouse brain using hybrid multiplexed sculpted light microscopy. Cell 177(4), 1050–1066 (2019)

[30] Klioutchnikov, A., Wallace, D.J., Frosz, M.H., Zeltner, R., Sawinski, J., Pawlak, V., Voit, K.-M., Russell, P.S.J., Kerr, J.N.: Three-photon head-mounted microscope for imaging deep cortical layers in freely moving rats. Nature methods 17(5), 509–513 (2020)

[31] Wang, M., Wu, C., Sinefeld, D., Li, B., Xia, F., Xu, C.: Comparing the effective attenuation lengths for long wavelength in vivo imaging of the mouse brain. Biomedical optics express 9(8), 3534–3543 (2018)

[32] Wang, T., Wu, C., Ouzounov, D.G., Gu, W., Xia, F., Kim, M., Yang, X., Warden, M.R., Xu, C.: Quantitative analysis of 1300-nm three-photon calcium imaging in the mouse brain. Elife 9, 53205 (2020)

[33] Mok, A.T., Wang, T., Zhao, S., Kolkman, K.E., Wu, D., Ouzounov, D.G., Seo, C., Wu, C., Fetcho, J.R., Xu, C.: A large field-of-view, single-cell-resolution two-and three-photon microscope for deep and wide imaging. eLight 4(1), 20 (2024)

[34] Podgorski, K., Ranganathan, G.: Brain heating induced by near-infrared lasers during multiphoton microscopy. Journal of neurophysiology 116(3), 1012–1023 (2016)

[35] Kolb, J.P., Klein, T., Kufner, C.L., Wieser, W., Neubauer, A.S., Huber, R.: Ultra-widefield retinal mhz-oct imaging with up to 100 degrees viewing angle. Biomedical Optics Express 6(5), 1534–1552 (2015)

[36] Mertz, J.: Introduction to Optical Microscopy. Cambridge University Press, ??? (2019)

[37] Beaulieu, D.R., Davison, I.G., Kilıç, K., Bifano, T.G., Mertz, J.: Simultaneous multiplane imaging with reverberation two-photon microscopy. Nature methods 17(3), 283–286 (2020)

[38] Wu, J., Liang, Y., Chen, S., Hsu, C.-L., Chavarha, M., Evans, S.W., Shi, D., Lin, M.Z., Tsia, K.K., Ji, N.: Kilohertz two-photon fluorescence microscopy imaging of neural activity in vivo. Nature methods 17(3), 287–290 (2020)

[39] Clough, M., Chen, I.A., Park, S.-W., Ahrens, A.M., Stirman, J.N., Smith, S.L., Chen, J.L.: Flexible simultaneous mesoscale two-photon imaging of neural activity at high speeds. Nature Communications 12(1), 6638 (2021)

[40] Li, X., Zhang, G., Wu, J., Zhang, Y., Zhao, Z., Lin, X., Qiao, H., Xie, H., Wang, H., Fang, L., et al.: Reinforcing neuron extraction and spike inference in calcium imaging using deep self-supervised denoising. Nature Methods 18(11), 1395–1400 (2021)

[41] Li, X., Li, Y., Zhou, Y., Wu, J., Zhao, Z., Fan, J., Deng, F., Wu, Z., Xiao, G., He, J., et al.: Real-time denoising enables high-sensitivity fluorescence time-lapse imaging beyond the shot-noise limit. Nature Biotechnology 41(2), 282–292 (2023)

[42] Wang, T., Xu, C.: Three-photon neuronal imaging in deep mouse brain. Optica 7(8), 947–960 (2020)

[43] Yao, Z., Velthoven, C.T., Kunst, M., Zhang, M., McMillen, D., Lee, C., Jung, W., Goldy, J., Abdelhak, A., Aitken, M., et al.: A high-resolution transcriptomic and spatial atlas of cell types in the whole mouse brain. Nature 624(7991), 317–332 (2023)

[44] Lau, C., Ng, L., Thompson, C., Pathak, S., Kuan, L., Jones, A., Hawrylycz, M.: Exploration and visualization of gene expression with neuroanatomy in the adult mouse brain. BMC bioinformatics 9, 1–11 (2008)

[45] Scott, B.B., Thiberge, S.Y., Guo, C., Tervo, D.G.R., Brody, C.D., Karpova, A.Y., Tank, D.W.: Imaging cortical dynamics in gcamp transgenic rats with a head-mounted widefield macroscope. Neuron 100(5), 1045–1058 (2018)

[46] Guo, C., Blair, G.J., Sehgal, M., Sangiuliano Jimka, F.N., Bellafard, A., Silva, A.J., Golshani, P., Basso, M.A., Blair, H.T., Aharoni, D.: Miniscope-lfov: A large-field-of-view, single-cell-resolution, miniature microscope for wired and wire-free imaging of neural dynamics in freely behaving animals. Science advances 9(16), 3918 (2023)

[47] Chen, Y., Jang, H., Spratt, P.W., Kosar, S., Taylor, D.E., Essner, R.A., Bai, L., Leib, D.E., Kuo, T.-W., Lin, Y.-C., et al.: Soma-targeted imaging of neural circuits by ribosome tethering. Neuron 107(3), 454–469 (2020)

[48] Grødem, S., Nymoen, I., Vatne, G.H., Rogge, F.S., Björnsdóttir, V., Lensjø, K.K., Fyhn, M.: An updated suite of viral vectors for in vivo calcium imaging using intracerebral and retro-orbital injections in male mice. Nature Communications 14(1), 608 (2023)

[49] Averbeck, B.B., Latham, P.E., Pouget, A.: Neural correlations, population coding and computation. Nature reviews neuroscience 7(5), 358–366 (2006)

[50] Pillow, J.W., Shlens, J., Paninski, L., Sher, A., Litke, A.M., Chichilnisky, E., Simoncelli, E.P.: Spatio-temporal correlations and visual signalling in a complete neuronal population. Nature 454(7207), 995–999 (2008)

[51] Stringer, C., Pachitariu, M., Steinmetz, N., Carandini, M., Harris, K.D.: High-dimensional geometry of population responses in visual cortex. Nature 571(7765), 361–365 (2019)

[52] Pinto, L., Rajan, K., DePasquale, B., Thiberge, S.Y., Tank, D.W., Brody, C.D.: Task-dependent changes in the large-scale dynamics and necessity of cortical regions. Neuron 104(4), 810–824 (2019)

[53] Bartolo, R., Saunders, R.C., Mitz, A.R., Averbeck, B.B.: Information-limiting correlations in large neural populations. Journal of Neuroscience 40(8), 1668–1678 (2020)

[54] Schultz, S.R., Copeland, C.S., Foust, A.J., Quicke, P., Schuck, R.: Advances in two-photon scanning and scanless microscopy technologies for functional neural circuit imaging. Proceedings of the IEEE 105(1), 139–157 (2016)

[55] Xiao, S., Cunningham, W.J., Kondabolu, K., Lowet, E., Moya, M.V., Mount, R.A., Ravasio, C., Bortz, E., Shaw, D., Economo, M.N., et al.: Large-scale deep tissue voltage imaging with targeted-illumination confocal microscopy. Nature Methods, 1–9 (2024)

[56] Kong, L., Cui, M.: A high throughput (> 90%), large compensation range, single-prism femtosecond pulse compressor. arXiv preprint arXiv:1306.5011 (2013)

[57] Pnevmatikakis, E.A., Giovannucci, A.: Normcorre: An online algorithm for piece-wise rigid motion correction of calcium imaging data. Journal of neuroscience methods 291, 83–94 (2017)

[58] Pnevmatikakis, E.A., Soudry, D., Gao, Y., Machado, T.A., Merel, J., Pfau, D., Reardon, T., Mu, Y., Lacefield, C., Yang, W., et al.: Simultaneous denoising, deconvolution, and demixing of calcium imaging data. Neuron 89(2), 285–299 (2016)

[59] Pachitariu, M., Stringer, C.: Cellpose 2.0: how to train your own model. Nature methods 19(12), 1634–1641 (2022)

[60] Kim, S.J., Slocum, A.H., Scott, B.B.: A miniature kinematic coupling device for mouse head fixation. Journal of Neuroscience Methods 372, 109549 (2022)

