## Supplementary Material for "Three-photon population imaging of subcortical brain regions"

### Three-photon population imaging of subcortical brain regions - Supplementary Information

### Contents

**Supplemental Note 1: Optimizing sampling rates for 3P calcium imaging of neuronal activity with GCaMP6s** 3

**Supplemental Note 2: Imaging improvement due to sampling optimization** 6

**Supplemental Note 3: The challenges in using commonly-used beam scanning mechanisms in large FOV 3P imaging** 10

**Extended Data Fig. 1: The fidelity of cell segmentation for sparse spatial sampling** 12

**Extended Data Fig. 2: The fidelity of calcium event extraction for sparse temporal sampling** 13

**Extended Data Fig. 3: Scan waveform and focal spot spacing** 14

**Extended Data Fig. 4: Heating simulation results** 15

**Extended Data Fig. 5: A comparison of denoising and filtering for postprocessing of datasets** 16

**Extended Data Fig. 6: A comparison of traces from denoised data and filtered data** 17

**Extended Data Fig. 7: 3P population imaging in the mouse hippocampus at video rate.** 18

**Extended Data Fig. 8: High resolution 3P imaging in the mouse hippocampus** 19

**Extended Data Fig. 9: The point-spread function of LIFT scope and the uniformity over its field of view** 20

**Extended Data Fig. 10: Field of view geometry** 21

**Supplemental Table 1: Acquisition parameters for all figures** 22

**Supplementary Video 1: Mouse brain structural stack** 23

**Supplementary Video 2: Large FOV 3P calcium imaging in mouse brain at around 1 mm depth** 24

### Supplemental Note 1: Optimizing sampling rates for 3P calcium imaging of neuronal activity with GCaMP6s

The desirable focal spot spacing for 3P calcium imaging of neuronal activity is the largest spacing that allows to spatially resolve cells and does not significantly affect the fidelity of cell segmentation. In order to find that spacing, we performed a systematic study of how focal spot spacing affects fidelity. First, we recorded a time series of calcium activity in a mouse hippocampus labeled with GCaMP6s, with small focal spot spacing (0.8 $\mu$m). The dataset was denoised, motion corrected and filtered to remove galvo-related artifacts, following our usual data analysis pipeline (see Methods). Then data points were removed to create additional time series with a gradually increasing focal spot spacing, from 1.6 $\mu$m to 10.4 $\mu$m (Fig. S1a-c). Those time series were then analyzed by our software that is based on CNMF. Due to the downsampling, cells appear to be different sizes in the different time series, necessitating a different parameter set as an input for CNMF for each one. To avoid extensive parameter sweeps within CNMF's large parameter space, which introduce variability and can skew results, we upsampled the resulting time series back to the native resolution using bicubic interpolation in Matlab before analyzing them with our software.

The result was a set of ROIs for each time series and the corresponding activity traces (Fig. S1d-f). To asses the fidelity of the ROI set retrieved for a particular large focal spot spacing, the correlation coefficient between each ROI retrieved from the large spacing set and all ROIs retrieved from the 0.8 $\mu$m spacing set was computed. For each ROI in the 0.8 $\mu$m dataset, the ROI from the large spacing set that had the highest correlation with it was chosen. A threshold of 0.6 correlation between the chosen ROI and the 0.8 $\mu$m dataset ROI was selected to determine whether the neuron was detected. This procedure was repeated for all ROIs in the 0.8 $\mu$m dataset. The true positive rate was computed as the number of detected cells in the large spacing set divided by the number of detected cells in the 0.8 $\mu$m data. False positive and false negative rates were computed in a similar manner, compared to the 0.8 $\mu$m data. Finally, the F-score [1] was computed from $\frac{TP}{TP+\frac{1}{2}(FN+FP)}$, where TP, FN and FP are the true positive rate, false negative rate and false positive rate, accordingly. The variance in F-score was computed by splitting the FOV of the original dataset into five and running this analysis pipeline on each of those datasets, in order to exclude variance associated with SNR, cell density, expression levels. etc.

The resulting plot shows a decreasing F-score with increasing focal spot spacing (Fig. 1g and S1), with the shaded region indicating the computed variance. However, the reduction in F-score from 0.8 $\mu$m to 4-5 $\mu$m focal spot spacing is minor, with the F-score only dropping by approximately 3% - less than the variance in cell segmentation resulting from the analysis with CNMF. The F-score starts to decline rapidly from 5-6 $\mu$m spacing and on, making 4 $\mu$m spacing optimal. Since the F-score of the 0.8 $\mu$m dataset is calculated against itself, the fit excludes that data point.

Similarly, the desirable frame rate for 3P calcium imaging with GCaMP6s is the minimal frame rate that allows to resolve calcium events. In order to find that frame rate, we performed a similar study of how the frame rate affects event extraction.

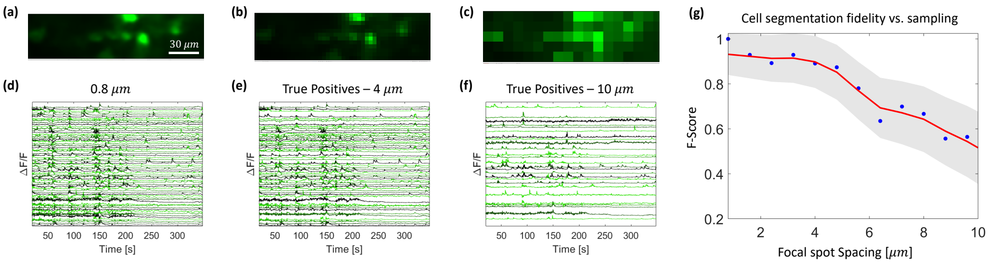


**Fig. S1** The fidelity of cell segmentation for sparse spatial sampling. (a)-(c) An example, zoomed-in section of the maximum intensity projection of the datasets with 0.8 $\mu$m, 4 $\mu$m, and 10 $\mu$m sampling, respectively. (d) The extracted traces for the 0.8 $\mu$m dataset. (e) The extracted traces from the 4 $\mu$m dataset, for the ROIs considered true positives when compared to the 0.8 $\mu$m dataset. (f) The extracted traces from the 10 $\mu$m dataset, for the ROIs considered true positives when compared to the 0.8 $\mu$m dataset. (g) Computed F-scores (blue dots) as a function of focal spot spacing, and a smoothing spline fit (red line) that serves as a guide to the eye. The shaded region indicates one standard deviation from the mean F-score.

First, we recorded a time series of neural activity in a mouse hippocampus labeled with GCaMP6s, with a 27 Hz frame rate. The dataset was denoised, motion corrected and filtered to remove galvo-related artifacts. Then frames were removed to create additional time series with a gradually decreasing frame rate, from 16 Hz to 0.25 Hz (Fig. S2a-c). Those time series were upsampled back to the native frame rate for comparison. All the resulting time series were then deconvolved to extract events using OASIS. The extracted event sequences were compared against those retrieved from the 27 Hz dataset, by computing the cosine similarity between the matrix of extracted events from all cells for the low frame rate dataset and the matrix of extracted events from all cells for the 27 Hz dataset. An event was considered detected if its timing was correct to within 0.2 s, half the rise time of GCaMP6s. The variance was computed by splitting the FOV of the original dataset into five and running this analysis pipeline on each of those datasets. The resulting plot shows an increasing similarity with increasing frame rate, with a minimal drop, from 1 to approximately 0.9, between 27 Hz to 6 Hz (Fig. S2d; error bars represents one standard deviation from the mean cosine similarity). We therefore chose 6 Hz as the design frame rate for LIFT.

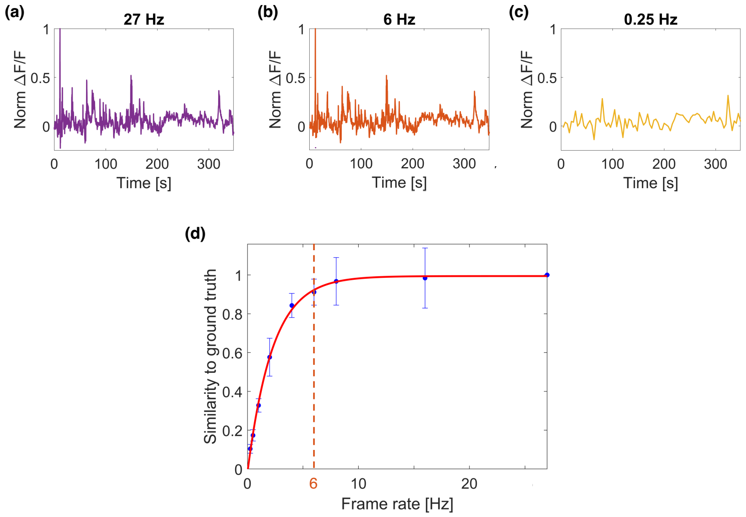


**Fig. S2** The fidelity of calcium event extraction as a function of frame rate. (a) An example trace recorded at a frame rate of 27 Hz. (b) The same trace as in b, downsampled to 6 Hz. (c) The same trace as in b, downsampled to 0.25 Hz. (d) Computed cosine similarities (blue dots) between the calcium events extracted from a dataset recorded at 27 Hz and temporally downsampled datasets, and an exponential fit (red line) to guide the eye. The error bars represents one standard deviation from the mean cosine similarity. The similarity between the calcium events extracted from the 27 Hz dataset and the calcium events extracted from the 6 Hz dataset, the design frame rate of LIFT, is $> 0.9$.

### Supplemental Note 2: Imaging improvement due to sampling optimization

Since tissue heating restricts the excitation power in 3P microscopy, optimized spatial and temporal sampling increases the SNR of calcium imaging data compared to conventional sampling for *any* FOV size, for two reasons. First, optimized spatial sampling allows to reduce the repetition rate of the source (or number of beamlets if using spatiotemporal multiplexing) because it uses fewer focal spots to sample the FOV and therefore requires a lower data acquisition rate. Similarly, optimized temporal sampling requires a lower acquisition rate because it uses less frames to sample an event. Reducing the repetition rate or beamlet number allows, in turn, for higher excitation pulse energies for the same excitation power. Due to the highly nonlinear dependence of the signal on pulse energy, higher pulse energies result in higher SNR, as explained below.

Secondly, with optimized spatial sampling, the excitation power can be increased beyond the levels used for small FOV imaging with conventional sampling, as the lower density of heat sources helps mitigate tissue heating. This further increases pulse energy and SNR. Overall, the SNR gain from sampling optimization can be utilized to improve FOV size, increase frame rate or to extend imaging depth.

When recording activity from a cell with a shot-noise limited system, the SNR of the recorded fluorescence trace scales with the number of spatial samples, $N_c$, within the cell body and the signal power per pixel, $S_{pp}$:

$$SNR \propto \sqrt{N_c S_{pp}} \tag{1}$$

For a single pulse per pixel measurement, the signal power per pixel depends on the energy of the pulse as $S_{pp} \propto {E_P}^3$, where $E_P$ is the pulse energy, and the pulse duration is fixed. Substituting that into 1 gives:

$$SNR \propto \sqrt{N_c {E_P}^3} \tag{2}$$

$N_c$, the number of spatial samples that fall within one cell, can be expressed as $N_c = A_c f_x^2$, where $A_c$ is the cell area, and $f_x = 1/x$ is the spatial sampling frequency, with $x$ being the distance between focal spots. For a given FOV size, the maximal $f_x$ possible depends on the frame rate, $f_{frame}$, and the repetition rate, $f_{rep}$, of the excitation source, assuming a single-pulse-per-pixel measurement:

$$N_c \propto f_x^2 \propto \frac{f_{rep}}{f_{frame}} = \frac{P_{exc}}{f_{frame} E_p} \tag{3}$$

Where $P_{exc}$ is the excitation power of the source, and $P_{exc} = E_P f_{rep}$. Since typically 3P systems use the maximum power that has been shown not to cause overheating for a given imaging depth, we will assume $P_{exc} = P_{max}$. Substituting $E_P \propto \frac{P_{max}}{N_c f_{frame}}$, into 2 gives:

$$SNR \propto \frac{1}{N_c} \left( \frac{P_{max}}{f_{frame}} \right)^{\frac{3}{2}} \tag{4}$$

Expressing this in terms of the spatial sampling frequency, $f_x$, gives:

$$SNR \propto \frac{1}{f_x^2} \left( \frac{P_{max}}{f_{frame}} \right)^{\frac{3}{2}} \tag{5}$$

Therefore, in principle the optimal sampling of a cell for a given excitation power is with a single spatial sample ($N_c$=1, $f_x \to 0$) and a single temporal sample ($f_{frame} \to 0$), as evident from Fig. 1i. Yet in practice there are several limitations. First, recording activity from cells labeled by an indicator requires a minimal frame rate in order to resolve that indicator's kinetics. For GCaMP6s, our data (Supplemental Note 1, Extended Data Fig. 2) shows that 6 Hz is the lowest frame rate that allows to maintain high calcium event extraction fidelity. Second, the number of spatial samples of a cell must yield the necessary spatial resolution to resolve it. For cells with an average diameter of 10 - 15 $\mu$m, our data (Supplemental Note 1, Extended Data Fig. 1) shows that 4 - 5 $\mu$m focal spot spacing is the largest spacing that preserves cell segmentation fidelity. We chose the average sampling distance to be 4 $\mu$m to account for the variation in focal spot spacing due to the nonlinearity of the scan (see Extended Data Fig. 3). In conventional 3P microscopy, focal spots are spaced by 0.5 - 1 $\mu$m. Using 1 $\mu$m sampling (Fig. S3a) a 9 x 11 $\mu$m cell is sampled by 70 - 80 focal spots, whereas in LIFT microscopy (Fig. S3b) it is sampled by 4 - 6 pixels. Consequently, the SNR of recorded fluorescence traces using LIFT microscopy is ~10 times higher than the SNR using 1 $\mu$m sampling of the same FOV, regardless of FOV size. The increase in SNR due to the use of optimized sampling allows to image deeper into the brain. Conversely, to achieve high enough SNR to image large FOVs at a 1 mm depth, conventional sampling schemes [2] would have to use an excitation power 10x higher than $P_{max}$, resulting in heating damage.

Moreover, due to the reduced density of heat sources, the excitation power in LIFT microscopy can be increased relative to the established maximum safe powers for conventional small FOV microscopy. To our knowledge, the safety of excitation powers in 3P microscopy has been assessed with a 0.5 $\mu$m sampling scheme over a 230 x 230 $\mu$m FOV [3]. Under those conditions, the maximum excitation power that was found to be safe was ~1 mW at the focal plane, for a depth of ~1 mm. Optimized spatial sampling enables higher excitation powers because the heat sources are spread over a larger area. Since the frame rates compatible with calcium imaging are fast enough to neglect tissue cooling between successive frames (~0.1°C/s) [3, 4], brain heating can be modeled as under continuous illumination of the focal spots of all the pixels in the FOV. Under those conditions, the total resulting temperature distribution is the sum of the temperature distributions from each focal spot, due to the linearity of the diffusion equation. As each focal spot will create a temperature increase over a certain local volume, the closer the focal spots are to each other, the more these volumes overlap. As a result, the total local temperature rise is lower for larger focal spot separations.

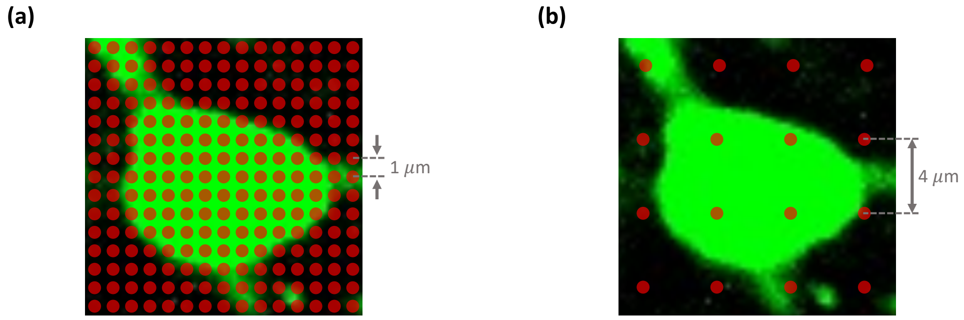


**Fig. S3** The sampling scheme used in LIFT microscopy. (a) An example of a conventional sampling scheme, with a 0.7 $\mu$m point-spread-function (PSF) and 1 $\mu$m focal spot spacing. A cell is sampled by 70 - 80 focal spots. (b) A schematic of the optimized sampling scheme used by LIFT, with a 0.7 $\mu$m PSF and 4 $\mu$m focal spot spacing. A cell is sampled by 4 - 6 focal spots.

To study the reduction in heating due to optimized sampling quantitatively, we numerically modeled the heating from a small FOV with 0.5 $\mu$m sampling and a large FOV with 4 $\mu$m sampling. Due to the high computational cost of a simulation with a spatial resolution that allows to resolve individual PSFs while simulating a centimeter-scale 3D volume, we performed two different numerical studies. The first, was adapted from Wang *et al.* [3] and previous work [1, 4], and simulated the entire volume (a cylinder of radius 6 mm and height 8 mm, as in [3]) but with uniform illumination over the entire FOV instead of individual Gaussian focal spots separated by the focal spot spacing. Brain heating was assessed using a Monte Carlo simulation to find the distribution of light scattered and absorbed, and then the finite difference method was used to evaluate the heating profile. We found that the brain heated more for a 230 $\mu$m FOV diameter (Fig. S4a) then for a 2 mm FOV diameter (Fig. S4b), and that the excitation power could be increased by a factor 2 for a 2 mm FOV diameter relative to the power for a 230 $\mu$m FOV diameter, while maintaining the same maximum temperature (Fig. S4c).

In another simulation, we studied the heating at the focal plane from individual focal spots that were either spaced by 0.5 $\mu$m or 4 $\mu$m (Fig. S4d-g). The simulation had a 2D circular geometry, and the simulated window extended to r = 400 $\mu$m, which was the maximum window size possible computationally with the resolution necessary to resolve the individual PSFs. We used a square array of 100 x 100 focal spots, and the temperature was fixed at 37°C at the simulated window boundary. In agreement with the uniform illumination simulation, the brain heated more for 0.5 $\mu$m focal spot spacing (Fig. S4d) than for 4 $\mu$m focal spot spacing (Fig. S4f) for 1 mW of power at the focal plane (Fig. S4e). In this simulation, the excitation power could be increased by a factor 3 for 4 $\mu$m focal spot spacing relative to 0.5 $\mu$m focal spot spacing, while maintaining the same maximum temperature (Fig. S4g). The possible power increase for 4 $\mu$m sampling could be larger in this simulation because it accounts for the reduced overlap between the heating profiles caused by individual focal spots and not

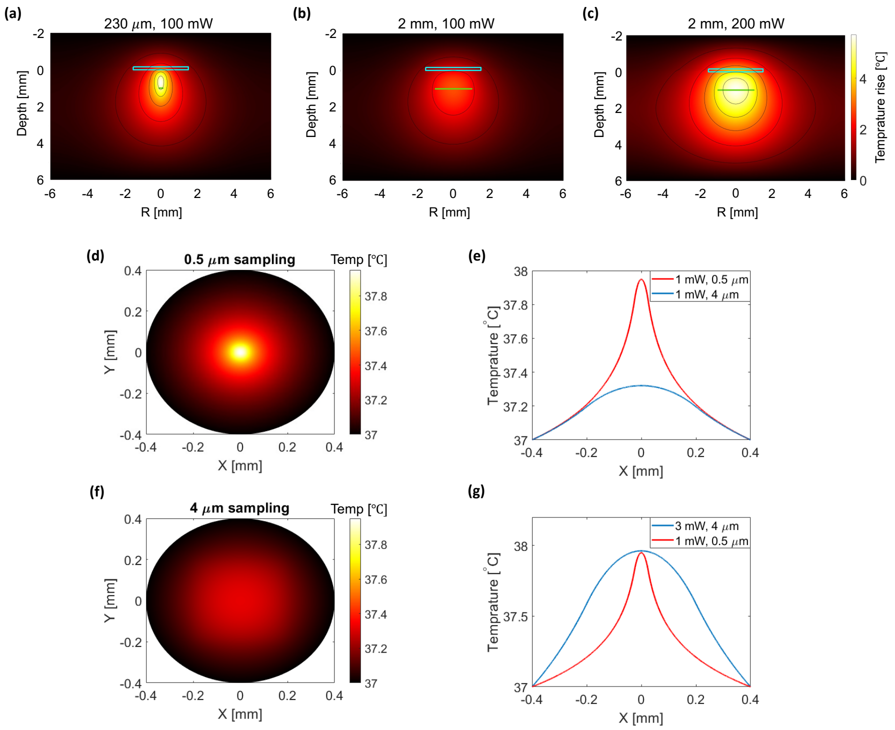


**Fig. S4** Heating simulation results. The temperature distribution resulting from imaging at a 1 mm depth with 1300 nm light in mouse brain, using uniform illumination over an area of diameter (a) 230 $\mu$m and 100 mW of power, (b) 2 mm and 100 mW of power, and (c) 2 mm and 200 mW of power. R represents the distance (radius) from center of the FOV and the color scale represents the increase in temperature during the imaging time and is the same for a-c. All powers are post objective. The green line indicates the scan area. The temperature distribution in the focal plane, resulting from continuous scanning of 100 x 100 focal spots with 1300 nm light and (d) 1 mW of excitation power with 0.5 $\mu$m sampling (e) 1 mW of excitation power with 4 $\mu$m sampling. (f) A comparison of the temperature distributions resulting from 1 mW excitation with 0.5 $\mu$m sampling and with 4 $\mu$m sampling, in the center of the focal plane (y=0). (g) A comparison of the temperature distributions resulting from 1 mW excitation with 0.5 $\mu$m sampling, and 3 mW excitation power with 4 $\mu$m sampling, in the center of the focal plane, showing that the maximum temperature in those two cases is similar.

just the reduced power density, which the uniform illumination simulation (Fig. S4a-c) accounts for. The gain in the SNR from increasing the excitation power can be utilized to improve imaging depth, or frame rate/FOV size by increasing the repetition rate.

### Supplemental Note 3: The challenges in using commonly-used beam scanning mechanisms in large FOV 3P imaging

As multiphoton microscopy is typically used to image inside scattering samples, spatial features cannot be resolved from a wide-field image of the emitted fluorescence. Rather, spatial resolution is commonly achieved by focusing the excitation light to a point and scanning the focus point by point over the sample plane. For each point, all the emitted fluorescence is collected and sent to a bucket detector, and the image is the composite two-dimensional array of detected intensities. In order to scan the beam, various scanning mechanisms are commonly used: a galvanometer, a resonant galvanometer, a polygon scanner and a micro-electro-mechanical system (MEMS) scanning mirror.

A galvanometer is the most commonly used beam scanning device. It allows for flexibility in the choice of scan speed, scan angle and scan waveform (i.e. the user can control the position of each individual scan point on the sample). Galvanometers have relatively large optical apertures and allow to scan large angles, albeit at compromised speeds. When using off-the-shelf optics, the beam is typically magnified from the galvanometer to the back aperture of the objective by a factor of $\sim 4$. With this geometry and a 1 MHz repetition rate, a high performance galvanometer can scan at about half the required speed for 4 $\mu$m sampling.

A resonant galvanometer is typically used for applications that require high-speed scanning. Large aperture resonant scanners can scan large angles at frequencies of several kHz. However, resonant scanners can typically scan only at their resonant frequency and only with a sinusoidal waveform. This property has two drawbacks: since the scan speed is set, so is the frame rate. Therefore, in one pulse per pixel imaging, the user can only choose pairs of FOVs and focal spot spacings that result in that frame rate. Furthermore, the sinusoidal scan pattern results in a large focal spot spacing in the center of the FOV and a small focal spot spacing at the edges. For a 2-mm-diameter FOV, a sinusoidal scan pattern that provides the desired frame rate of 6 Hz would result in a focal spot spacing of $> 6$ $\mu$m at the center of the FOV. Finally, resonant scanner frequency can vary based on the particular mirror geometry and often cannot be controlled to provide the exact desired frequency and thus the desired focal spot spacing.

A polygon scanner can scan large angles with large optical apertures, and its scan speed can be tuned. Since beam scanning is based on rotation of the polygon, it is the only scanner that does not oscillate back and forth and therefore is the only scanner that can provide a completely linear scan and uniform sampling. However, the scan angle cannot be varied and thus neither can the FOV. Furthermore, polygon scanners with optical apertures that are large enough to fill the back aperture of a typical objective with a 4x magnification as described above, require very large scan angles. These scan angles typically require specially designed, bulky scan lenses. Finally, polygon scanners suffer from pupil shifting due to the shifting of the location of the reflection point from the scanner facet during rotation.

A MEMS scanner allows for both large scan angles and large scan speeds, and provides the ability to tailor the scan waveform. Yet currently MEMS scanners that support those scan speeds and angles have relatively small optical apertures. Using such apertures would either result in significant underfilling of the objective back aperture, or in a reduction of the scanned FOV due to a magnification larger than 4x.

Our custom scanning module overcomes all of these challenges by using a galvanometer relay, as described in Fig. 2a. The use of a galvanometer relay doubles the scan speed of a galvanometer while preserving all of its advantages, allowing the user to choose any combination of frame rate, focal spot spacing and FOV possible given the laser repetition rate. The scan waveform can be tailored for large FOV calcium imaging so that the focal spot spacing in the FOV center does not significantly exceed 5 $\mu$m, yet the overall scan frequency results in the desired frame rates. Furthermore, our scanner enables random access scanning, which is used in a variety of applications, such as targeted illumination microscopy [5] and patterned optogenetic stimulation [6]. Finally, our scanner can be easily replicated, using all off-the-shelf components.

### Extended Data Fig. 1: The fidelity of cell segmentation for sparse spatial sampling

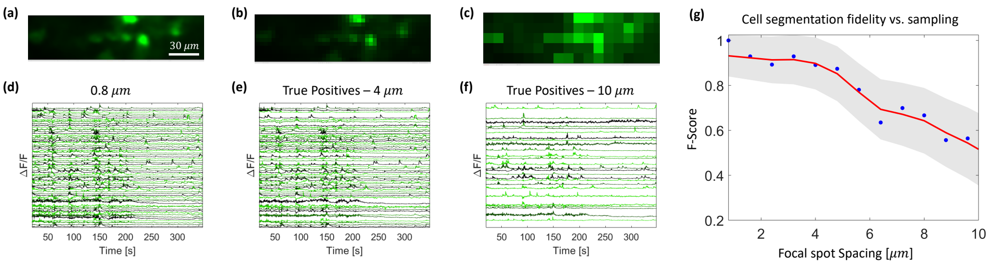


Calculated cell segmentation fidelity, following the procedure detailed in Supplemental Note 1. (a)-(c) An example, zoomed-in section of the maximum intensity projection of the datasets with 0.8 $\mu$m, 4 $\mu$m, and 10 $\mu$m sampling, respectively. (d) The extracted traces for the 0.8 $\mu$m dataset. (e) The extracted traces from the 4 $\mu$m dataset, for the ROIs considered true positives when compared to the 0.8 $\mu$m dataset. (f) The extracted traces from the 10 $\mu$m dataset, for the ROIs considered true positives when compared to the 0.8 $\mu$m dataset. (g) Computed F-scores (blue dots) as a function of focal spot spacing, and a smoothing spline fit (red line) that serves as a guide to the eye. The shaded region indicates one standard deviation from the mean F-score.

### Extended Data Fig. 2: The fidelity of calcium event extraction for sparse temporal sampling

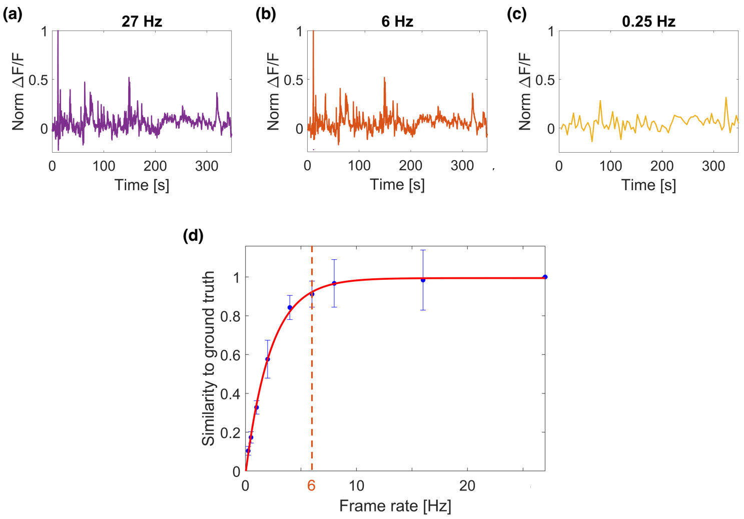


Calculated event extraction fidelity, following the procedure detailed in Supplemental Note 1. (a) An example trace recorded at a frame rate of 27 Hz. (b) The same trace as in b, downsampled to 6 Hz. (c) The same trace as in b, downsampled to 0.25 Hz. (d) Computed cosine similarities between the calcium events extracted from a dataset recorded with 27 Hz and temporally downsampled datasets. The error bars represents one standard deviation from the mean, and the red line is an exponential fit. The similarity between the event sequences extracted from the 27 Hz dataset and the event sequences extracted from the 6 Hz dataset, the design frame rate of LIFT, is $> 0.9$.

### Extended Data Fig. 3: Scan waveform and focal spot spacing

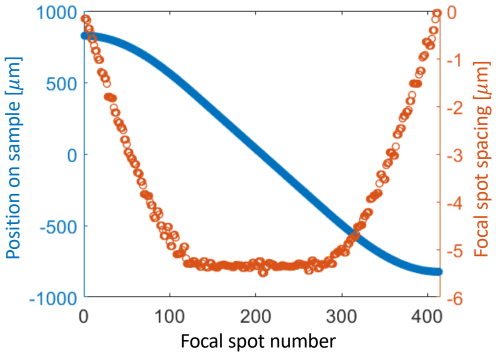


The lateral position of the beam in the fast axis of the sample plane as a function of focal spot number (or time), for the waveform used for a 3 mm$^2$ FOV scan at a 6 Hz frame rate (blue circles), and the corresponding spacing between adjacent focal spots (orange circles).

### Extended Data Fig. 4: Heating simulation results

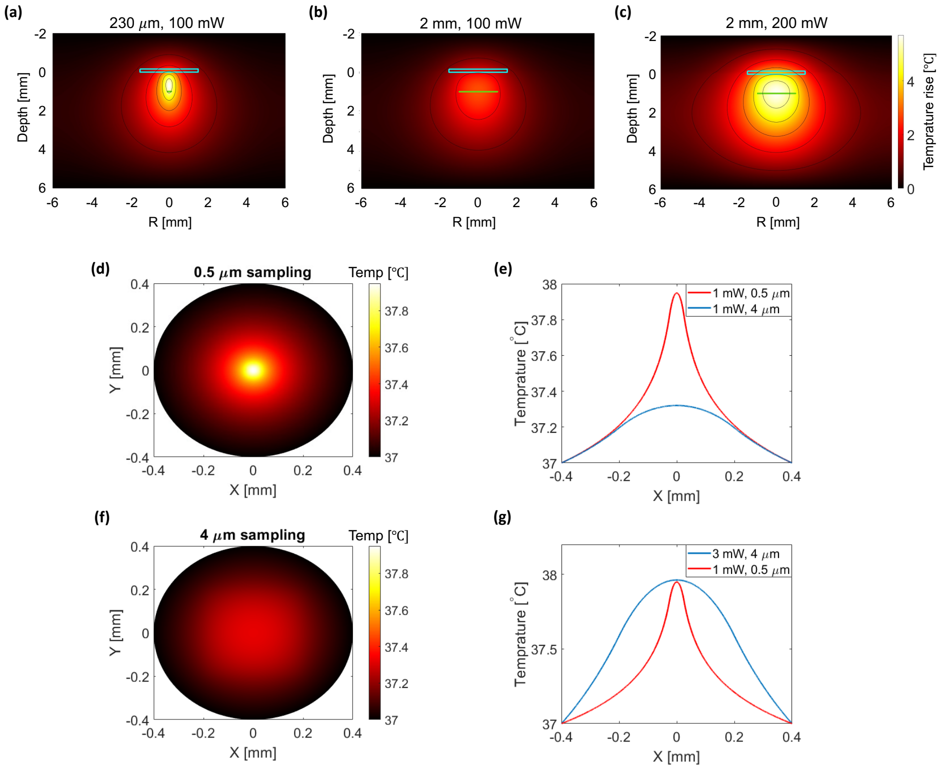


Results of brain tissue heating simulations, detailed in Supplemental Note 2. The temperature distribution resulting from imaging at a 1 mm depth with 1300 nm light in mouse brain, using a uniform illumination over an area of diameter (a) 230 $\mu$m and 100 mW of power, (b) 2 mm and 100 mW of power, and (c) 2 mm and 200 mW of power. R represents the distance (radius) from center of the FOV and the color scale represents the increase in temperature during the imaging time and is the same for a-c. All powers are post objective. The green line indicates the scan area. The temperature distribution in the focal plane, resulting from continuous scanning of 100 x 100 focal spots with 1300 nm light and (d) 1 mW of excitation power with 0.5 $\mu$m sampling (e) 1 mW of excitation power with 4 $\mu$m sampling. (f) A comparison of the temperature distributions resulting from 1 mW excitation with 0.5 $\mu$m sampling and with 4 $\mu$m sampling, in the center of the focal plane (y=0). (g) A comparison of the temperature distributions resulting from 1 mW excitation with 0.5 $\mu$m sampling, and 3 mW excitation power with 4 $\mu$m sampling, in the center of the focal plane, showing that the maximum temperature in those two cases is similar.

### Extended Data Fig. 5: A comparison of denoising and filtering for postprocessing of datasets

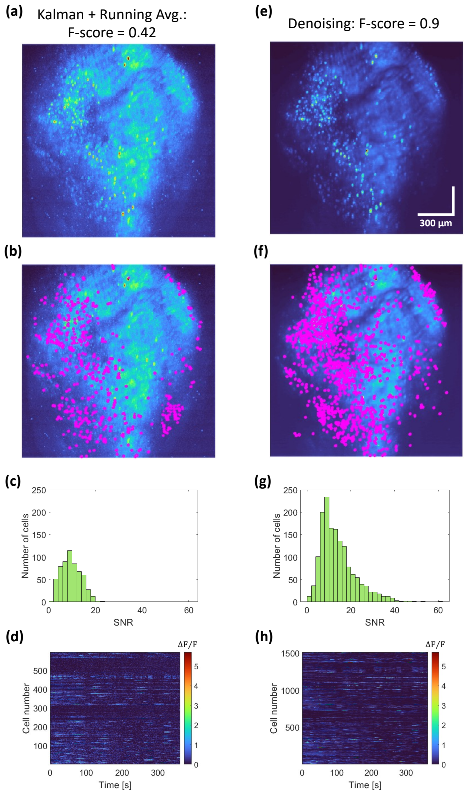


A comparison of the dataset analyzed in Fig. 4a-e, processed by filtering (a-d) and processed by denoising (e-h). (a) The maximum intensity projection (MIP) of the filtered time series. The data was processed using our data processing pipeline, yet instead of denoising, the time series was filtered with a Kalman filter with 0.75 gain and then smoothed with a temporal running average of length 6. (b) Contours of 570 active cells, plotted in magenta, on top of the MIP. The fidelity of cell segmentation, evaluated using the F-score (see Supplemental Note 1), was low (F-score = 0.42). (c) A histogram of the SNRs of the extracted activity traces. For the comparison, the noise was defined as the standard deviation of the top 20% of frequencies and the signal was defined as in the main text. (d) The full data set, including traces from all active cells. The color scale corresponds to the normalized $\Delta F/F$ value. (e)-(h) The same as a-d, but for the denoised data. The number of cells detected in the denoised data is nearly triple the number detected in the filtered data, and the fidelity of cell segmentation is more than double (F-score = 0.9). For a direct comparison of the extracted traces, see Extended data Fig. 6.

### Extended Data Fig. 6: A comparison of traces from denoised data and filtered data

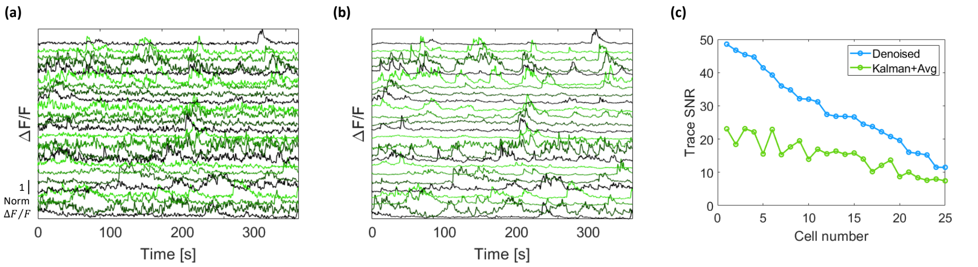


(a) Traces retrieved from 25 example ROIs from the data presented in Extended data Fig. 5a-d, which was filtered with a Kalman filter with 0.75 gain and then smoothed with a temporal running average of length 6. (b) Traces retrieved from the same ROIs as in a, but from the denoised dataset, presented in Extended data Fig. 5e-h. (c) A comparison of the SNRs of the traces in a and b. The cells are numbered according to their order in figures a-b, with Cell 1 being the top trace. The SNR was defined as in Extended data Fig. 5. Overall, the SNR of the denoised data is higher while the calcium events are preserved.

### Extended Data Fig. 7: 3P population imaging in the mouse hippocampus at video rate.

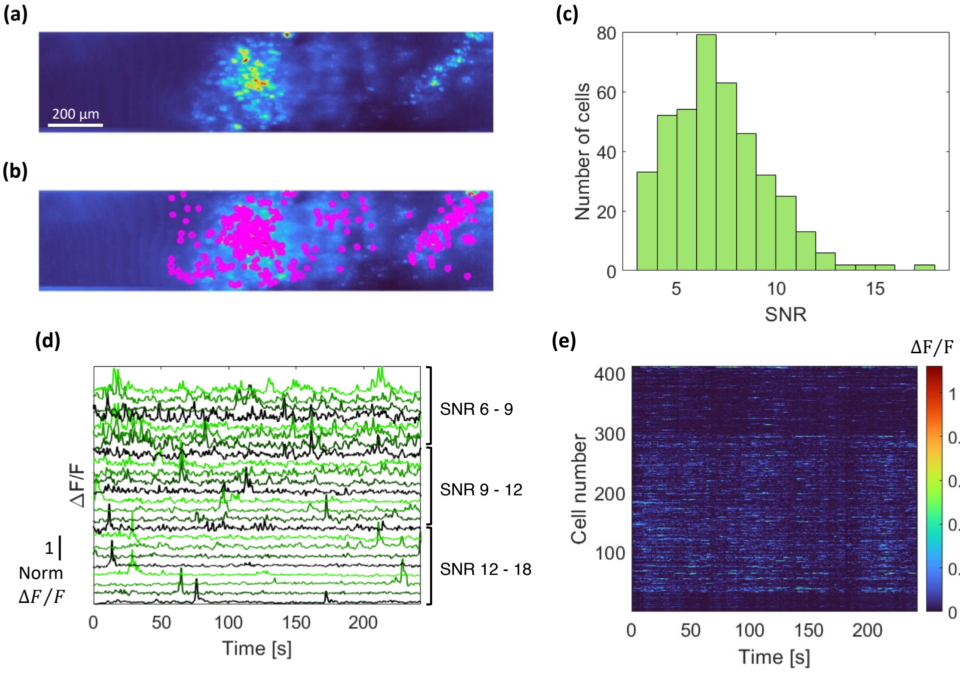


Neuronal activity recording at a 27 Hz frame rate, over a 0.73 mm$^2$ FOV, at a depth of 900 $\mu$m in the mouse hippocampus. (a) The maximum intensity projection (MIP) of the time series. (b) Contours of 400 active cells, plotted in magenta, on top of the MIP. (c) A histogram of the SNRs of the extracted activity traces shows that the majority of cells have 3 $<$ SNR $<$ 8. (d) Examples of traces from various SNR ranges. (e) The full data set, including traces from all active cells. Imaging was performed with 4 $\mu$m lateral sampling and cells were labeled with GCaMP6s.

### Extended Data Fig. 8: High resolution 3P imaging in the mouse hippocampus

**(a)** **(b)**

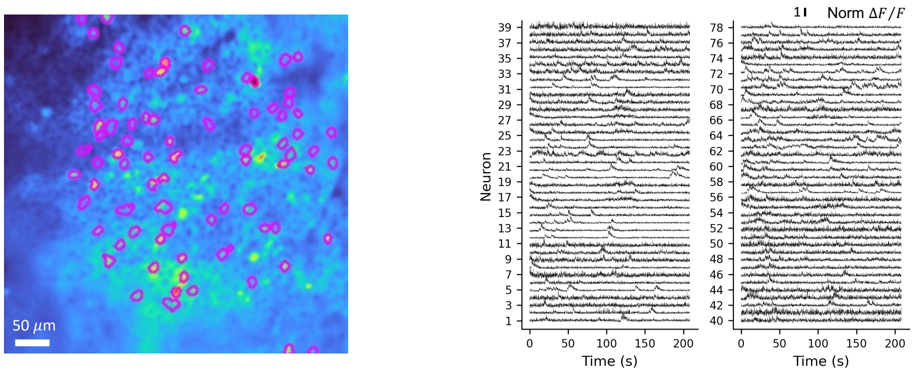


Neuronal activity recording with 2 $\mu$m lateral sampling at a 10 Hz frame rate, over a 500 x 500 $\mu$m FOV, at a depth of 948 $\mu$m in the mouse hippocampus. The data was motion corrected and artifacts were removed but it was not denoised or filtered. (a) The mean intensity projection of the recorded time series. Contours of active cells segmented using our CNMF-based software are plotted on top in magenta. (b) The corresponding activity traces, normalized to the maximum value of the trace.

### Extended Data Fig. 9: The point-spread function of LIFT scope and the uniformity over its field of view

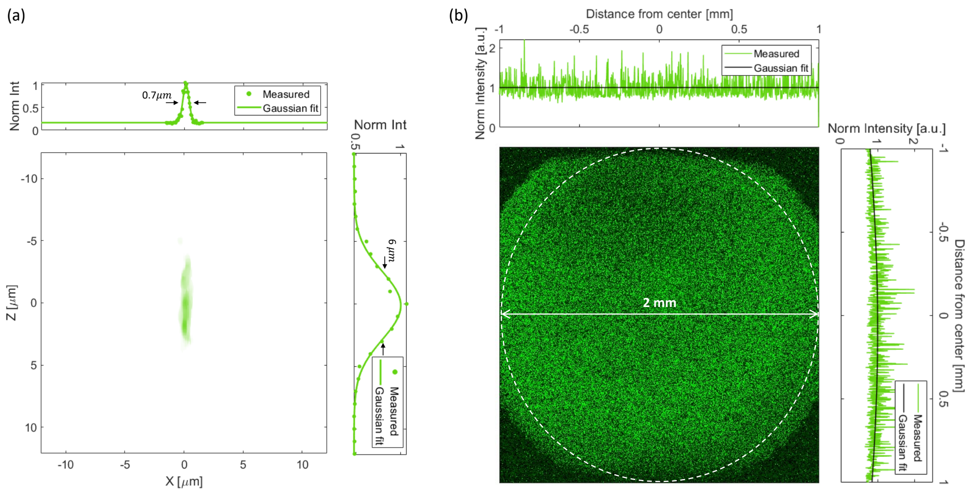


(a) The 3P point spread function (PSF) of LIFT scope, measured on 0.5 $\mu$m fluorescent beads. Top: The intensity profile of the measured lateral PSF (green dots) and a Gaussian fit with a FWHM of 0.7 $\mu$m (green line). Right: The intensity profile of the measured axial PSF (green dots) and a Gaussian fit with a FWHM of 6 $\mu$m (green line). Some waviness is noticeable in the z-axis of the PSF image, due to a slight wobble in the mechanical scan. (b) The measured 3P signal from a uniform sample (fluorescein solution) over the FOV of LIFT scope. The white dashed circle represents LIFT's full, 2-mm-diameter FOV. Top: The intensity profile across a horizontal line through the center (green line) and a Gaussian fit (black line). Right: The intensity profile across a vertical line through the center (green line) and a Gaussian fit (black line).

### Extended Data Fig. 10: Field of view geometry

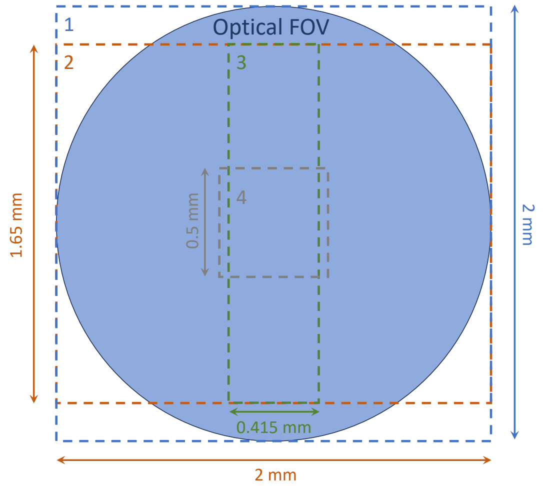


A schematic showing the scanned FOV for the datasets shown in this work. 1 - Scanned FOV for high resolution structural imaging, presented in Fig. 3 and 6. 2 - Scanned FOV for 6 Hz calcium imaging, presented in Fig. 4 and 5. 3 - Scanned FOV for video-rate calcium imaging, presented in Extended data Fig. 7. 4 - Scanned FOV for high resolution 10 Hz calcium imaging, presented in Extended data Fig. 8.

### Supplemental Table 1: Acquisition parameters for all figures

| Figure | Animal | Dwell time (μs) | Excitation Power (mW) | Avg. focal spot spacing (μm) | Frame rate (Hz) | Frames averaged | FOV geometry | Imaging type |
|---|---|---|---|---|---|---|---|---|
| 3 | Mouse | 4 | 10-160 | 1 | 0.06 | 1-10 | 1 | Structural |
| 4, 5, ED 5, ED 6 | Mouse | 1 | 107-180 | 4 | 6 | 1 | 2 | Activity |
| 6 | Rat | 4 | 50-120 | 1 | 0.06 | 3-10 | 1 | Structural |
| ED 7 | Mouse | 1 | 150 | 4 | 27 | 1 | 3 | Activity |
| ED 8 | Mouse | 1 | 107 | 2 | 11 | 1 | 4 | Activity |

Acquisition parameters for all main text and extended date figures. ED denotes extended data figures, the FOV geometries refer to the schematics shown in Extended Data Fig. 10.

#### Supplementary Video 1: Mouse brain structural stack

A structural stack in the mouse brain with a 2 mm diameter FOV and 1 $\mu$m lateral sampling (see Fig. 3). The images were taken at 250 - 1230 $\mu$m depth with a 5 $\mu$m step.

#### Supplementary Video 2: Large FOV 3P calcium imaging in mouse brain at around 1 mm depth

A denoised large FOV time series recorded in the mouse brain at a depth of 981 $\mu$m, with 4 $\mu$m lateral sampling and a 6 Hz frame rate. The analyzed data is presented in Fig. 4. The frame rate of the video was increased to 30 Hz for clarity.

### References


[1] Demas, J., Manley, J., Tejera, F., Barber, K., Kim, H., Traub, F.M., Chen, B., Vaziri, A.: High-speed, cortex-wide volumetric recording of neuroactivity at cellular resolution using light beads microscopy. Nature Methods **18**(9), 1103–1111 (2021)

[2] Mok, A.T., Wang, T., Zhao, S., Kolkman, K.E., Wu, D., Ouzounov, D.G., Seo, C., Wu, C., Fetcho, J.R., Xu, C.: A large field-of-view, single-cell-resolution two-and three-photon microscope for deep and wide imaging. eLight **4**(1), 20 (2024)

[3] Wang, T., Wu, C., Ouzounov, D.G., Gu, W., Xia, F., Kim, M., Yang, X., Warden, M.R., Xu, C.: Quantitative analysis of 1300-nm three-photon calcium imaging in the mouse brain. Elife **9**, 53205 (2020)

[4] Podgorski, K., Ranganathan, G.: Brain heating induced by near-infrared lasers during multiphoton microscopy. Journal of neurophysiology **116**(3), 1012–1023 (2016)

[5] Demmerle, J., Innocent, C., North, A.J., Ball, G., Müller, M., Miron, E., Matsuda, A., Dobbie, I.M., Markaki, Y., Schermelleh, L.: Strategic and practical guidelines for successful structured illumination microscopy. Nature protocols **12**(5), 988–1010 (2017)

[6] Oron, D., Papagiakoumou, E., Anselmi, F., Emiliani, V.: Two-photon optogenetics. Progress in brain research **196**, 119–143 (2012)
